# BRUTUS-LIKE proteins moderate the transcriptional response to iron deficiency in roots

**DOI:** 10.1101/231365

**Authors:** Jorge Rodríguez-Celma, Robert T. Green, James M. Connorton, Inga Kruse, Yan Cui, Hong Qing Ling, Janneke Balk

## Abstract

Iron is an essential micronutrient but in excess is toxic inside cells. Under iron deficiency, the expression of iron uptake genes is increased, but it is not known how the transcriptional response is controlled to avoid uptake of too much iron. The hemerythrin E3 ligases BRUTUS (BTS) and BTS-LIKE (BTSL) have previously been identified as negative regulators of the iron deficiency response. Our phylogenetic analysis indicated that BTSL proteins are present in dicotyledonous plants only and form a separate clade from BTS homologs. *BTSL1* and *BTSL2* in *Arabidopsis thaliana* are in a network with nearly all iron uptake genes, whereas *BTS* is in a shoot-specific network. *BTSL1* and *BTSL2* are expressed predominantly in the root epidermis and cortex, separate from *BTS* in the root stele, shoot and embryos. Mutant analysis identified *BTSL2* as the dominant paralog of the otherwise redundant *BTSL* genes. The *btsl* double mutant had increased protein levels of FIT, the FER-like Iron deficiency-induced Transcription factor, and failed to switch off the transcriptional response upon iron resupply, leading to dramatic iron accumulation in roots and shoots. Protein interaction between the C-terminus of BTSL proteins and FIT indicate that FIT is a direct target for degradation. Taken together, our studies show that *BTSL1* and *BTSL2* control iron uptake in the epidermis and cortex, upstream of BTS in the vasculature and leaves.

## INTRODUCTION

Iron (Fe) is the fourth most abundant element in the Earth’s crust, but its bioavailability is greatly limited by the insolubility of Fe hydroxides. Plants have developed highly efficient Fe uptake systems, classically divided in Strategy I or reductive strategy, present in dicotyledonous plants, and Strategy II or chelating strategy in grasses (Römheld and Marschner, 1986; Tsai and Schmidt, 2017). The reductive strategy in plants such as Arabidopsis involves the activity of a Ferric Reductive Oxidase, FRO2, to reduce Fe^3+^ to Fe^2+^ which is then taken up by the Iron-Regulated Transporter IRT1. A key regulator of Fe uptake in Strategy I plants is the basic Helix-Loop-Helix (bHLH) transcription factor FIT (FER-like Iron deficiency-induced Transcription factor; Bauer et al., 2007). FIT forms heterodimers with one of four bHLH proteins from subgroup Ib, namely bHLH38, bHLH39, bHLH100 and bHLH101. Mutant studies combined with transcriptomics have identified more than 400 genes that are controlled by FIT (Colangelo and Guerinot, 2004; Mai et al., 2016). These targets include *FRO2* and *IRT1* for which direct promoter binding by the FIT-bHLH Ib dimer has been shown (Yuan et al., 2008; Wang et al., 2013).

How the Fe status of the plant is communicated to the transcriptional regulators remains unknown. Recent studies have discovered a small family of hemerythrin E3 ligases in plants, including BRUTUS (BTS) in Arabidopsis (Long et al., 2010; Selote et al., 2015) and Hemerythrin motif-containing RING- and Zn-finger proteins (HRZ1, HRZ2) in rice (Kobayashi et al., 2013), that act as negative regulators of the Fe deficiency response. The expression of *HRZ1, HRZ2* and *BTS* is induced ~10-fold under Fe deficiency. Mutant lines of *BTS* or *HRZ* showed increased tolerance to Fe deficiency and accumulation of Fe in leaves and seeds. BTS / HRZ proteins contain three putative Fe-binding hemerythrin domains in the N-terminus, and a CHY/RING-type Zn finger domain with ubiquitin E3 ligase activity in the C-terminus (Kobayashi et al., 2013; Selote et al., 2015). This domain architecture bears similarity to mammalian FBXL5 (F-box/LRR-repeat protein 5), which targets the Iron Regulatory Protein 2 (IRP2) for degradation (Vashisht et al., 2009; Salahudeen et al., 2009). The activity of FBXL5 depends on Fe binding to its N-terminal hemerythrin domain and on oxygen levels (Salahudeen et al., 2009). Plants do not have a functional homolog of IRP2, however, BTS was shown to interact with, and affect the stability of bHLH105 (ILR3) and bHLH115, two Fe-regulated transcription factors (Selote et al., 2015). A heterodimer of ILR3 and bHLH104 regulates the expression of *bHLH38* / *39* / *100* / *101,* encoding the main protein partners of FIT (Zhang et al., 2016).

*BTS* was first identified as *EMBRYO DEFECTIVE 2454* (*EMB2454*) in a screen for embryo lethal mutants (McElver et al., 2001; Tzafrir et al., 2004), and later re-named and described as a negative regulator of Fe homeostasis (Long et al., 2010). Since *bts* loss-of-function mutants are lethal, studies have focused on viable alleles with T-DNA insertion in the promoter/UTRs or point mutations (Selote et al., 2015; Hindt et al., 2017). These *bts* mutant plants showed an enhanced Fe deficiency response, including increased root length, soil acidification and higher Fe reductase activities, and they are more tolerant to Fe limiting conditions. Promoter-GUS lines showed that *BTS* is expressed in leaves and embryos independent of the Fe status of the plant, and its expression is induced in the stele of the root under Fe deficiency (Selote et al., 2015). BTS is not the only hemerythrin-E3 ligase in Arabidopsis, as previously noticed by Kobayashi and colleagues (2013). Two homologous genes are found in the Arabidopsis genome, *BRUTUS-Like1* (*AT1G74770*) and *BRUTUS-Like2* (*AT1G18910*). Double *btsl1 btsl2* mutants have recently been reported to be more tolerant to Fe deficiency and to accumulate more Fe (Hindt et al., 2017). A triple mutant of *btsl1, btsl2* and a viable *bts* allele had an enhanced phenotype, and the authors concluded that the *BTSL* genes act as redundant negative regulators of Fe homeostasis together with *BTS* (Hindt et al., 2017).

Here we show that the function of the *BTSL* genes differs from *BTS. BTSL1 and BTSL2* are expressed predominantly in the root epidermis and cortex and are part of a root-specific transcriptional network of genes involved in Fe uptake. In contrast, *BTS* is co-expressed with only a small number of shoot-specific Fe-regulated genes. We found that *BTSL1/2* function to control the levels of FIT protein and prevent excess Fe accumulation via the FRO2 / IRT1 uptake pathway.

## RESULTS

### *BTSL1* and *BTSL2* are co-expressed with Fe-responsive genes in the roots

Co-expression analysis is a powerful tool for functional characterization because co-regulated genes are likely to function in the same biochemical pathway or biological process (Stuart et al., 2003). Previously we surveyed the transcriptional response to Fe deficiency in Arabidopsis and in *Medicago truncatula* (Rodríguez-Celma et al., 2013a). We considered the overlap between the two data sets working under the assumption that Fe homeostasis regulation should be conserved between plant species. We found the orthologous pair *AT1G18910-Medtr7g037040* up-regulated under Fe deficiency with the core genes of the Fe regulon. This gene has recently been annotated as *BTSL2* (Hindt et al., 2017). As an independent and unbiased approach, we used publicly available microarray data to build co-expression networks around *BTSL2* and the two homologous genes *BTSL1* and *BTS.* First, we found that *BTSL1* and *BTSL2* are co-regulated with each other but not with *BTS.* Second, we noticed that a large network could be built around *BTSL1* and *BTSL2* with root-specific data sets, but no co-regulated ‘root’ partners could be found for *BTS* (Supplementary Figure S1 and Supplementary Table S1). Correlations with *BTS* were only found when using shoot-specific data sets. Because *BTS, BTSL1* and *BTSL2* have been linked to the Fe deficiency response, the networks were trimmed to only those genes present in the so-called ferrome (Schmidt and Buckhout, 2011) (Figure 1A, B, Supplementary Table S2). We found that *BTSL1* and *BTSL2* are co-regulated with genes involved in Fe uptake, such as the Fe transporter *IRT1* and genes for the biosynthesis and export of coumarin-derived Fe chelators *(PDR9, 4CL2* and *F6H1)* (Figure 1A). Also co-expressed with *BTSL1* and *BTSL2* are the transcription factor *FIT* and its partner *bHLH39,* central regulators in the Fe-deficiency response (Figure 1B). Basically, the entire root Fe uptake regulon is found, except for *bHLH38* and *FRO2,* but these genes are not present on the micro-array chip (ATH1) that is most commonly used in the public database. The *BTS* network includes the transcription factor *POPEYE* (*PYE*), as previously documented (Long et al., 2010), and the ferric reductase *FRO3.* Also found is *AT5G05250,* a gene of unknown function which is highly induced during Fe deficiency (Rodriguez-Celma et al., 2013b).

**Figure 1.**
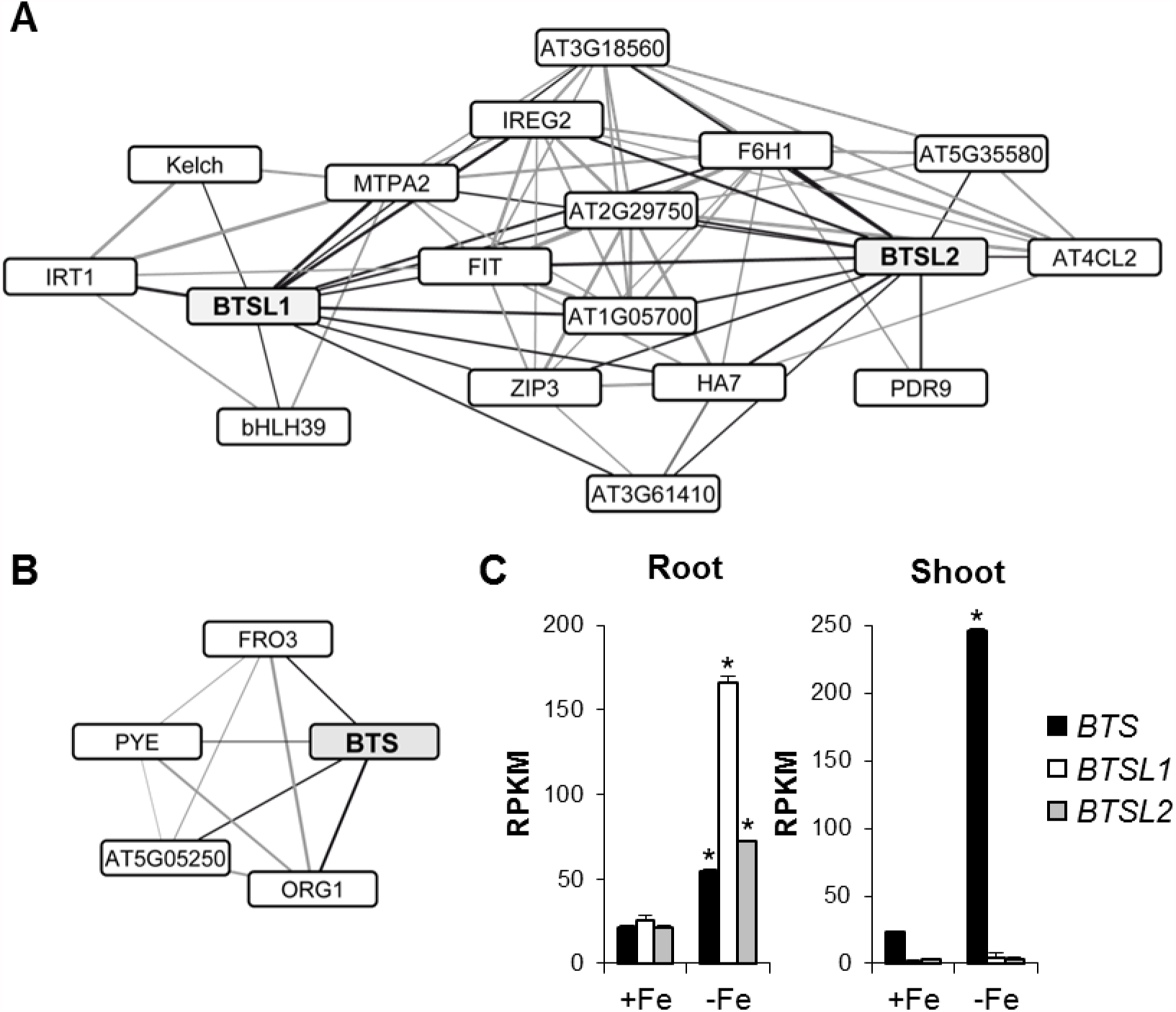
*BTSL* and *BTS* are part of different gene expression networks. **(A, B)** Co-expression analysis of Fe-responsive genes that are co-regulated with (A) *BTSL1* and *BTSL2* in roots, and (B) *BTS* in shoots. The edge thickness relate to the Pearson’s correlation coefficient. See Supplemental Table S2 for descriptions of all genes and numerical values. **(C)** Expression levels of *BTS, BTSL1* and *BTSL2* in roots and in shoots under Fe sufficient (+Fe) and Fe deficient conditions (-Fe). Data were obtained by RNAseq as described in Rodriguez-Celma et al., (2013). RPKM, Reads Per Kilobase Million. The bars represent the mean of n = 3 biological replicates x 10 seedlings ± SD (* p<0.05 using a two-tailed *t*-test).

Investigation of the expression of *BTS, BTSL1* and *BTSL2* in RNAseq data sets of the Fe deficiency response (Rodríguez-Celma et al., 2013b) confirmed that *BTSL1* and *BTSL2* were expressed mainly in the roots and strongly induced under Fe deficiency (Figure 1C). *BTS* is therefore the dominant family member expressed in the shoots, in accordance with the calculated networks.

### BTSL1 and BTSL2 proteins are specific to dicotyledonous plants

To study the evolutionary relationship between BTS and BTSL, we performed a phylogenetic analysis of homologous protein sequences across the tree of life. As noted previously (Kobayashi et al., 2013; Matthiadis & Long, 2016) homologs containing more than one hemerythrin (Hr) domain fused to a RING-domain E3 ligase are found only among plant species, including mosses. Interestingly, Arabidopsis BTSL1 and BTSL2 are in a separate clade unique to dicotyledonous species (Figure 2A).

**Figure 2.**
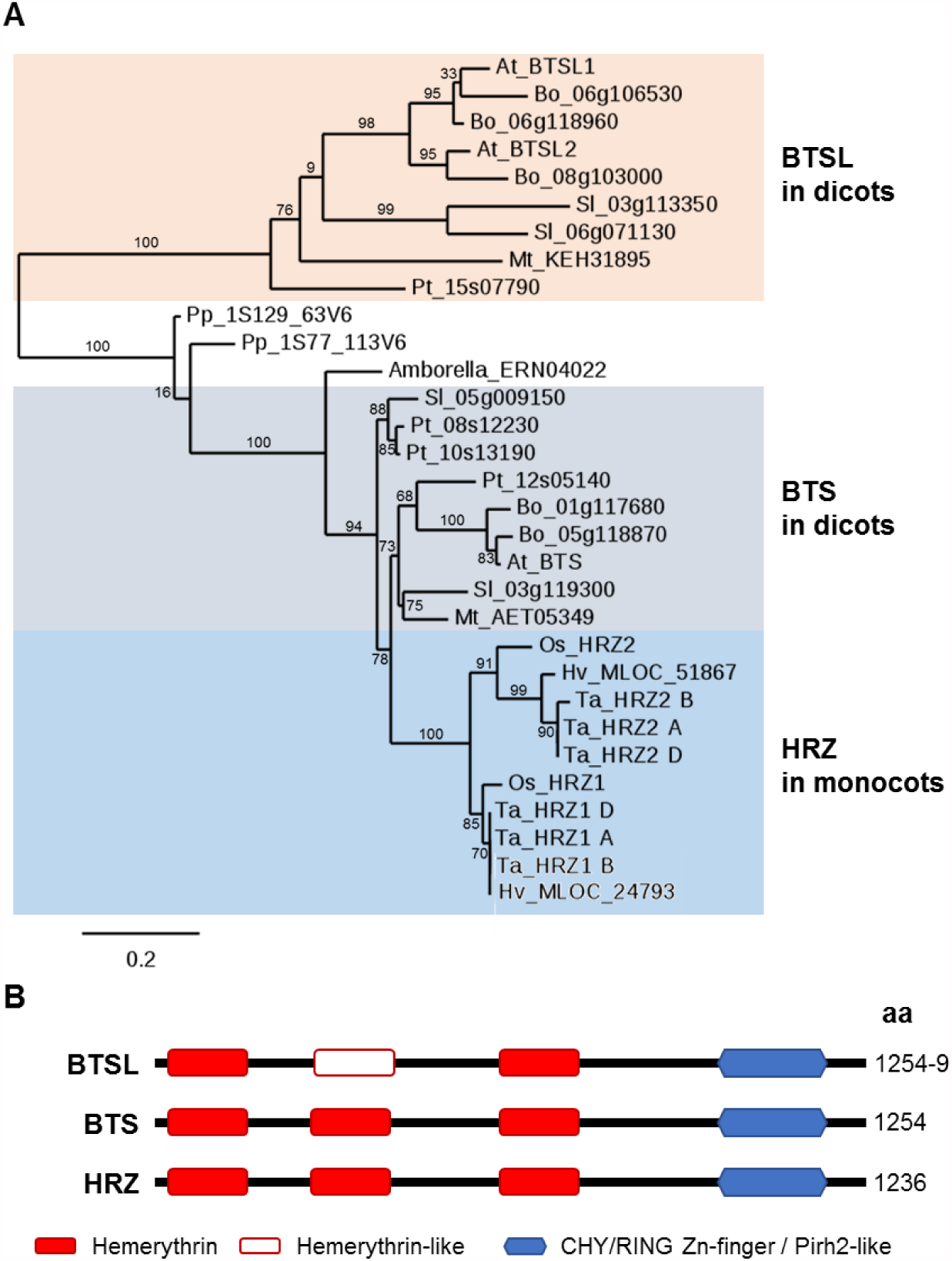
BTSL are uniquely found in dicotyledonous plants. **(A)** Phylogenetic tree of BTS homologs from selected plant species. Sequences were found by BLASTing the amino acid sequences of Arabidopsis BTS, BTSL1, BTSL2 and rice HRZ1 in Ensembl Plants (http://plants.ensembl.org). Species used: Amborella, *Amborella trichocarpa;* At, *Arabidopsis thaliana;* Bo, *Brassica oleracea;* Hv, *Hordeum vulgare;* Mt, *Medicago truncatula;* Os, *Oryza sativa;* Pp, *Physcomitrella patens;* Pt, *Populus trichocarpa;* Sl, *Solanum lycopersicum;* Ta, *Triticum aestivum.* Numbers next to branches indicate bootstrapping values for 100 replications. The scale bar indicates the rate of evolutionary change expressed as number of amino acid substitutions per site. **(B)** Domain organization of BTSL, BTS and HRZ proteins.

BTSL1/2 proteins have two predicted Hr domains and a CHY/RING Zn-finger domain (Kobayashi et al., 2013). The two Hr domains of BTSL1/2 correspond to the first and third Hr domain in BTS. Further inspection of the amino acid sequence of BTSL1 and 2 revealed a third putative Hr domain (Figure 2B), not recognized by domain searching algorithms. Alignment of the individual Hr domains of the three proteins showed that in the gap between the two Hr domains of BTSL1 and 2 the amino acid sequence is highly homologous to the other Hr sequences, but lacks the four conserved histidine residues found in the canonical Hr domains. This domain is still predicted to form a four α-helical bundle typical of hemerythrins (Supplementary Figure S2) and additional histidines and acidic residues are present which may function as ligands for Fe. The C-terminal Zn finger domain is highly conserved, with 80% amino acid identity between BTSL1 and BTSL2, and 65% identity with BTS. This domain consists of a unique arrangement of Zn-fingers, a CHY/RING-type Zn finger domain found in the mammalian E3 ligase Pirh2 / RCHY1 (Leng et al., 2003) and in four other E3 ligases in Arabidopsis (Lee and Seo, 2016).

### *BTSL1* and *BTSL2* promoters are activated by Fe deficiency in specific root tissues

To investigate where in the roots and in which cell types *BTSL1* and *BTSL2* are expressed, the promoter regions of these genes were cloned into the vector pBGWFS7 upstream of an *eGFP-GUS* fusion reporter gene (Karimi et al., 2002). For the *BTSL1* promoter we used the intergenic region of 880 nt upstream of the ATG start codon. In the case of *BTSL2* the intergenic region was ~5 kb, and therefore we decided to use only 2097 nt upstream of the ATG start codon. Seeds were germinated on agar plates with minimum salts (see materials and methods section) supplemented either with 50 μM FeEDTA (+Fe) or with 100 μM ferrozine to effectively deplete Fe (-Fe). After 5 days of growth seedlings were analysed for GFP expression. Aside from some autofluorescence in the empty seed coat, no GFP fluorescence was observed in any part of the seedling under Fe sufficient conditions (+Fe). However, under Fe deficiency (-Fe), GFP was highly induced in the roots (Figure 3A), in agreement with RNAseq expression data (Figure 1C). At this early growth stage, GFP expression was similar in the *BTSL1* and *BTSL2* promoter constructs. In older seedlings that had developed lateral roots, the expression pattern of the *BTSL1* and *BTSL2* promoters became different: GUS staining showed that *BTSL1* is expressed in the upper half of the root under Fe deficiency, whereas *BTSL2* is expressed in the lower half of the root, predominantly in the differentiation zone (Figure 3B). *BTSL1* and *BTSL2* promoter-GUS activity was also found in the root cap (columella), but not in the root meristem or elongation zone. Cross-sections of the GUS-stained roots showed that both promoters are induced in the epidermis, including root hairs, and cortex cells. *BTSL2* was also expressed in the endodermis and stele in the differentiation zone (Figure 3B). The expression patterns of *BTSL1* and *BTSL2* contrast with the pattern observed for *BTS. BTS* promoter-GUS expression showed little promoter activity in the epidermis and cortex, but was strongly upregulated in the stele in response to Fe deficiency (Selote et al., 2015). Moreover, *BTS* is expressed in embryos which lack *BTSL1* and *BTSL2* promoter activity (Supplementary Figure S3).

**Figure 3.**
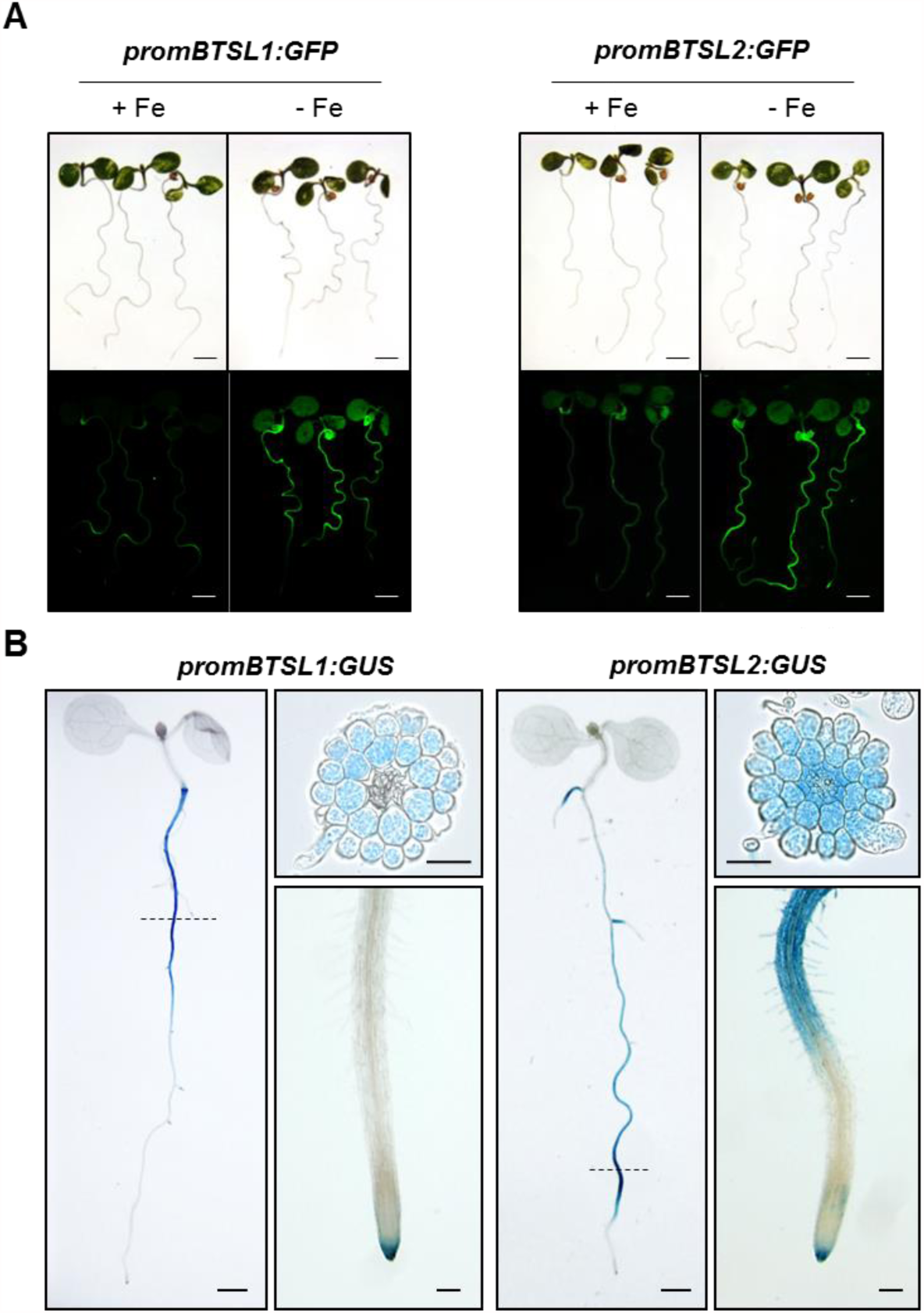
Promoter activity of *BTSL1* and *BTSL2* in Fe-deficient roots. Promoter sequences of *BTSL1* (-880 nt) and *BTSL2* (-2097 nt) were inserted upstream of eGFP-GUS and the constructs were stably expressed in wild-type Arabidopsis. **(A)** GFP fluorescence in seedlings grown on medium with 50 μM FeEDTA (+Fe) or with 100 μM ferrozine to deplete Fe (-Fe), as indicated, for 6 days. Scale bar is 1 mm. **(B)** GUS activity staining in seedlings grown on medium depleted of Fe (100 μM ferrozine) for 8 days. Scale bar is 0.5 mm for whole seedlings, 100 μm for a close-up of the root tip, and 50 μm for root cross sections. Images are representative of 3 independent lines for each promoter construct.

In summary, the *BTSL1* and *BTSL2* promoters are activated by Fe deficiency in the upper and lower part of the root, respectively. Both genes are expressed in epidermis, cortex and columella cells.

### *BTSL1* and *BTSL2* have partially redundant functions in maintaining Fe homeostasis

To characterise the function of *BTSL* in Fe homeostasis, T-DNA insertion lines were obtained from the Nottingham Arabidopsis Stock Centre. Three independent mutant lines were selected for *BTSL1* (SALK_015054, SALK_032464 and SALK_117926; named *btsl1-1, btsl1-2* and *btsl1-3,* respectively), but only one line with a T-DNA insertion in the coding sequence was available for *BTSL2* (SALK_048470; *btsl2-2)* (Figure 4A). The *btsl1-1* was described as *btsl1* in Hindt et al., (2017) and the *btsl2* allele used by the same authors has an insertion in the promoter. Homozygous lines were selected by PCR, T-DNA insertion sites were confirmed by sequencing, and transcript levels were assessed by RT-qPCR. The expression of *BTSL1* in roots under standard and Fe deficient conditions was virtually abolished in the *btsl1-1* and *btsl1-3* alleles, but residual expression remained in *btsl1-2* (Figure 4B). Disruption of the *BTSL1* gene led to a 2-fold upregulation of *BTSL2* expression under Fe deficiency (Figure 4C). The *btsl2-2* mutant lacked detectable levels of *BTSL2* transcript (Figure 4C), while *BTSL1* expression was ~3-fold higher than wild type lacking Fe (Figure 4B). Double knock-out lines were produced by crossing *btsl1-1* and *btsl1-3* with the *btsl2-2* line. The *btsl1-1 btsl2-2* mutant was genetically complemented by either *BTSL1* or *BTSL2* (see below).

**Figure 4.**
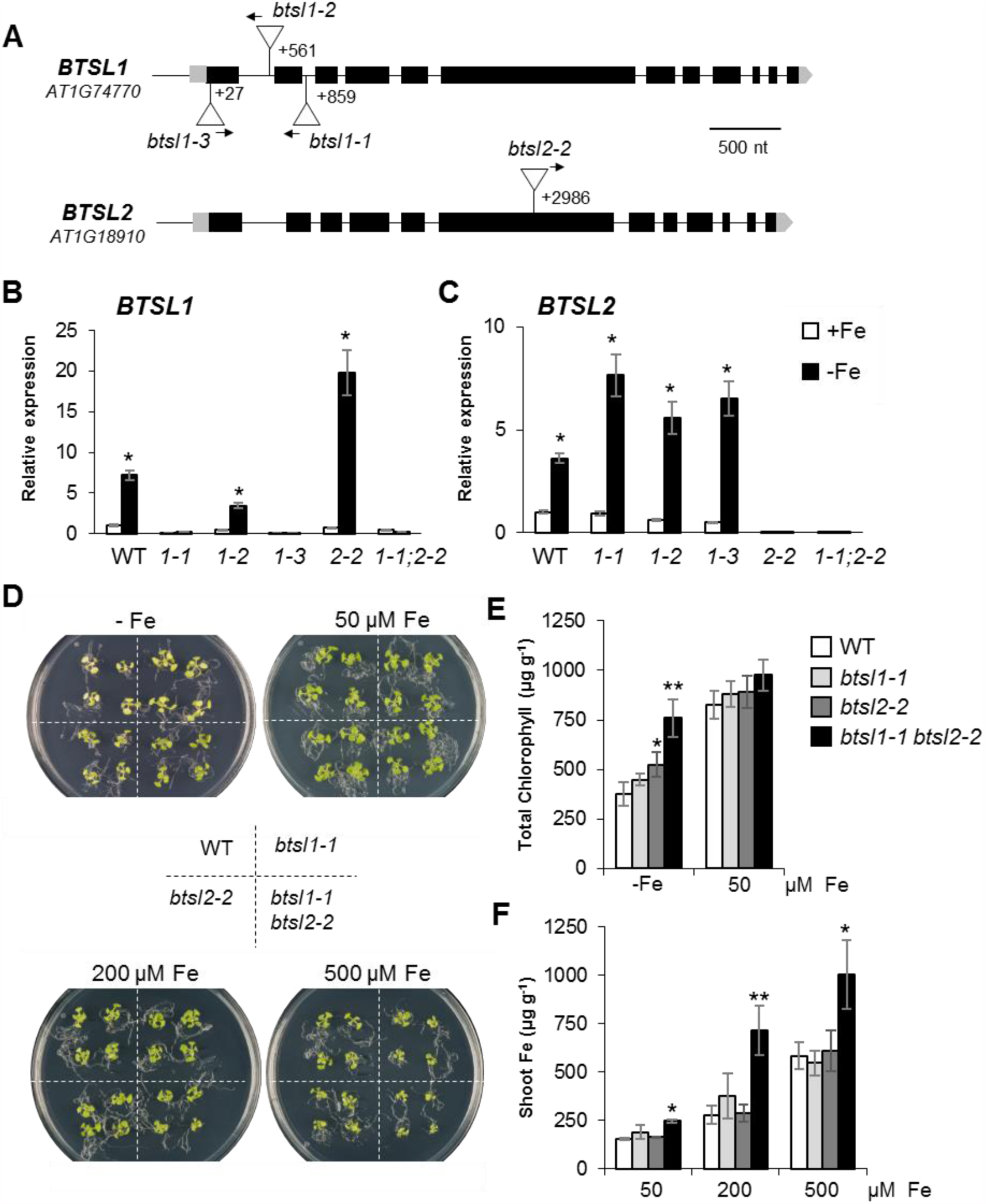
*BTSL1* and *BTSL2* have redundant functions in Fe homeostasis. **(A)** Schematic representation of the *BTSL1* and *BTSL2* genes and mutant alleles. Exons are in black, 5’ and 3’-UTRs in grey. T-DNA insertions are indicated by triangles: *btsl1-1,* SALK_015054; *btsl1-2,* SALK_032464; *btsl1-3,* SALK_117926; *btsl2-2,* SALK_048470. Arrows indicate the left border primer used for genotyping. **(B, C)** Transcript levels of *BTSL1* (B) and *BTSL2* (C) in wild-type and mutant alleles under standard (+Fe, 50 μM FeEDTA) and Fe-deficient conditions (-Fe, 100 μM ferrozine). (* p<0.05 using a two-tailed t-test). **(D)** Growth of wild-type and *btsl* alleles on medium with varying Fe concentrations. Seeds were germinated on medium with 50 μM FeEDTA, and, after 10 days, transferred to medium depleted of Fe using 100 μM ferrozine (-Fe) or 50 μM FeEDTA; or grown in the presence of 200 or 500 μM FeEDTA for 14 days. **(E, F)** Chlorophyll concentration (E) under -Fe and control conditions, and iron concentration in shoots (F) from wild type (WT) and *btsl* mutant alleles, grown under control and Fe toxicity conditions as in (D). The bars represent the mean of n = 3 biological replicates x 10 seedlings ± SD (* p<0.05; ** p<0.01 using a two-tailed *t*-test).

Mutant lines were tested for phenotypes on a range of Fe concentrations. Seedlings were germinated and grown on medium with 50 μM FeEDTA for 10 days, followed by 4 days on plates with ferrozine (-Fe), or 4 days on Fe sufficient medium (50 μM FeEDTA). The effect of toxic concentrations of Fe was tested by growing the seedlings on plates with 200 or 500 μM FeEDTA for 14 days. Single insertion lines showed no obvious phenotype under any of the conditions tested. On the other hand, the *btsl1-1 btsl2-2* double mutant line showed subtle but noticeable phenotypes under conditions of both Fe deficiency and Fe excess (Figure 4D). After 4 days without Fe, the *btsl* double mutant appeared less chlorotic than wild type, which was confirmed by chlorophyll measurements (Figure 4E). The *btsl2-2* mutant allele also retained some chlorophyll, suggesting that *BTSL1* and *BTSL2* are not fully redundant. In the presence of excess Fe, growth of the *btsl* double mutant was significantly more impaired than wild type or the single mutants (Figure 4D; Supplementary Figure S4). Quantification of Fe levels in shoots showed a 2-fold accumulation of this metal in *btsl* plants grown on excess Fe (Figure 4F), indicating that this might be the cause of the observed growth impairment.

### *BTSL1* and *BTSL2* limit Fe uptake after a period of Fe deficiency

Because *BTSL1* and *BTSL2* are only expressed in Fe-deficient roots, but phenotypes were seen under Fe excess, we tested whether the genes have a specific function in the transition from Fe deficiency to Fe sufficiency. Wild-type and *btsl* double mutant seedlings were grown up on medium with Fe (50 μM FeEDTA) for ten days and then transferred to medium without Fe but with ferrozine for 3 days to induce the Fe deficiency response. After that, plants were resupplied with Fe by transferring them back to Fe sufficient conditions, and then sampled after 3 days. In the control treatment, seedlings were always grown on 50 μM Fe for 10 + 3 + 3 days, with transfers to fresh medium. A diagram of the treatments and time points of sampling is presented in Figure 5A. For comparison, we also grew the *brutus-1* (*bts-1*) mutant line (Selote et al., 2015) under the same conditions. *bts-1* is a viable allele that lacks mRNA induction under Fe deficiency.

**Figure 5.**
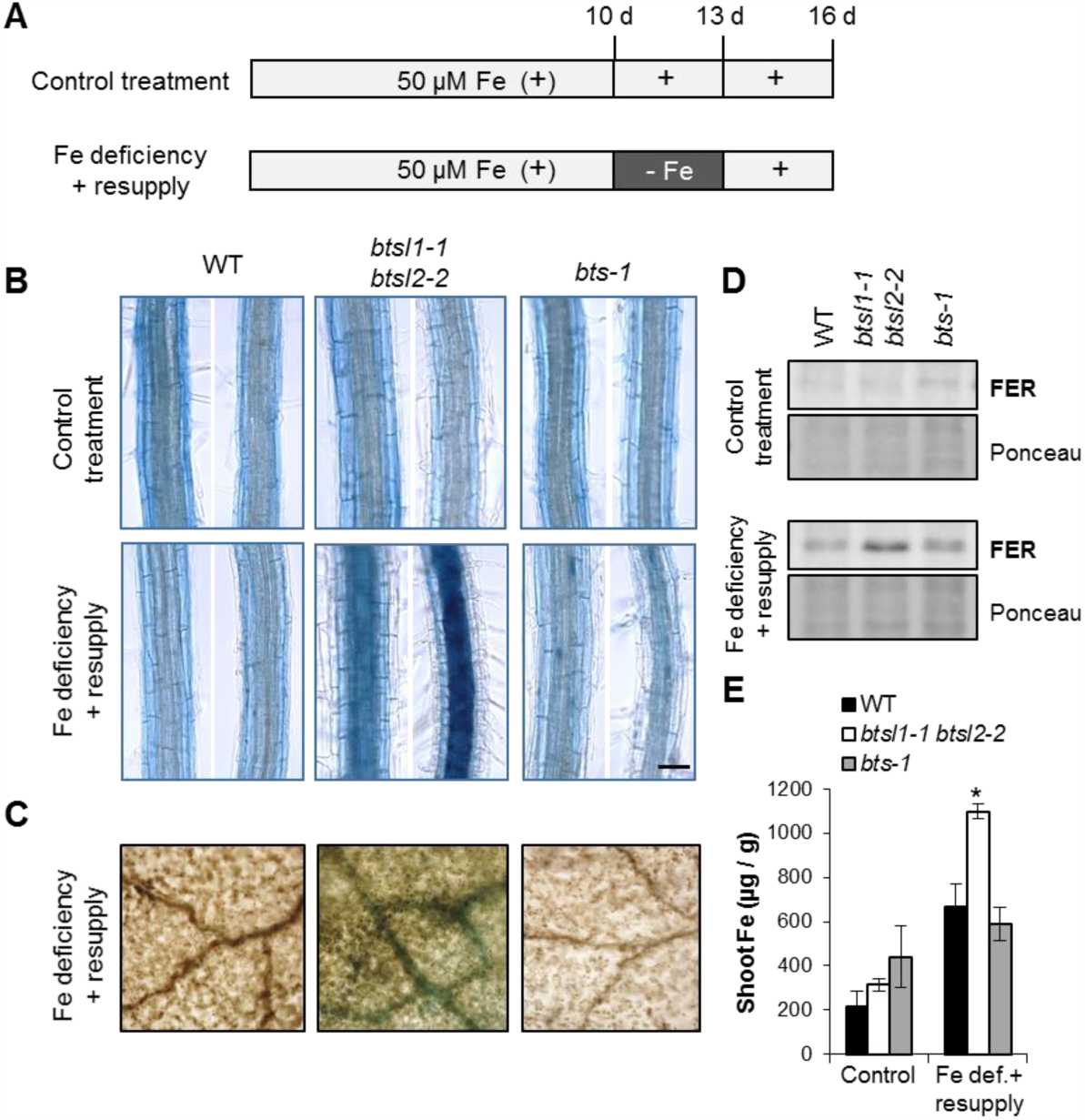
The *btsl* double mutant hyper-accumulates Fe after a period of deficiency. **(A)** Diagram of the Fe treatments used in this study. Seedlings were germinated on agar plates with 50 μM FeEDTA. On day 10, seedlings were transferred to a new plate (control treatment) or to medium depleted of Fe (100 μM ferrozine). After 3 days, seedlings were transferred back to medium with 50 μM FeEDTA. Samples were taken at the end of the treatment. **(B)** Perls’ Prussian Blue staining for Fe in roots in wild-type, *btsl1-1 btsl2-2* double mutant and *bts-1* seedlings following control (top) and Fe deficiency-resupply treatments (bottom) according to the diagram in (A). The images show a section of the differentiation zone. Scale bar is 50 μm. **(C)** Perls’ Prussian Blue staining of leaves after the Fe deficiency-resupply treatment. The images are a close-up of the adaxial leaf surface with veins. Scale bar is 75 μm. **(D)** Immunoblot of ferritin (FER) protein levels in root samples of wild type, *btsl1-1 btsl2-2* double mutant and *bts-1* seedlings after control and resupply treatments. **(E)** Quantification of Fe in the shoots of wild-type, *btsl1-1 btsl2-2* double mutant and *bts-1* seedlings after control and Fe deficiency-resupply treatments. Bars represent the mean of 3 biological replicates of 10 pooled plants each ± SD (* p<0.05 using a two-tailed t-test).

None of the genotypes showed chlorosis or growth impairment after the Fe deficiency and resupply treatment. However, the *btsl* double mutant accumulated large amounts of Fe in the central core of the root and in leaf veins as shown by Perls’ staining (Figure 5B, C). Iron did not accumulate when mutants were grown continuously on 50 μM Fe. *bts-1* mutants behaved as wild-type plants and did not show increased Perls’ staining in any part of the root or leaves after the Fe deficiency-resupply treatment.

To gain further insight into the management of the Fe influx, we analysed the levels of ferritin protein, which functions as a temporary Fe store in Arabidopsis roots and leaves (Ravet et al., 2009). Ferritin protein levels followed the same trend as observed with Fe staining, with ferritin accumulating only in the *btsl* double mutant after the Fe deficiency-resupply treatment (Figure 5D).

When we measured the Fe content of the shoots, we found that wild-type plants, when subjected to deficiency and resupply, contained 3-fold more Fe compared to continuous growth on 50 μM Fe (Figure 5E). It is likely that upregulation of the Fe uptake machinery during Fe deficiency leads to a sudden influx of Fe when this is resupplied. In wild-type seedlings, the increased Fe levels do not seem to be toxic as there is no growth impairment and Fe is evenly distributed (Figure 5B, C). In the *btsl* double mutant, the Fe content of shoots increased more than 5-fold after the deficiency-resupply treatment relative to the control conditions (Figure 5E), reaching similar levels to those observed in plants grown continuously with 500 μM Fe (Figure 4F). In contrast to the *btsl* double mutant, *bts-1* seedlings had a similar Fe content to wild type after the Fe deficiency-resupply treatment (Figure 5E).

In summary, our data show that the *btsl* double mutant is unable to adapt to changes in Fe supply, accumulating toxic concentrations of Fe. Moreover, the lack of Fe accumulation in *bts-1* line under these conditions suggests that BTSLs and BTS have different functions or act in different tissues.

### The *btsl1-1 btsl2-2* double mutant fails to switch off the Fe deficiency response

Next, we investigated the levels of FRO2 and IRT1, two key players in Fe uptake, during Fe deficiency and resupply in the wild-type and *btsl* double mutant. We followed the same treatment described above, and plants were sampled before the resupply from the control (+Fe) and Fe deficiency treatment (-Fe), and daily after Fe resupply (1, 2 and 3 d Fe resupply) (Figure 6A). Wild-type seedlings were able to regulate FRO2 enzyme activity as expected, showing a ~6-fold induction in response to Fe deficiency and returning down to basal levels one day after Fe resupply (Figure 6A). In *btsl* double mutant seedlings FRO2 enzyme activity remained high after Fe resupply: the activity was 4-fold induced after one day, 3-fold induced after two days and it was still nearly double the basal wild-type levels after three days (Figure 6A). The single insertion *btsl1-1* and *bts-1* mutant lines behaved like the wild type, but *btsl2-2* showed a slight delay in switching off FRO2 activity (Supplementary Figure S5A). The sustained induction of FRO2 activity in the *btsl* double mutant was restored by expressing either *35S:BTSL-YFP* or *35S:BTSL2-GFP* (Supplementary Figure S5B).

**Figure 6.**
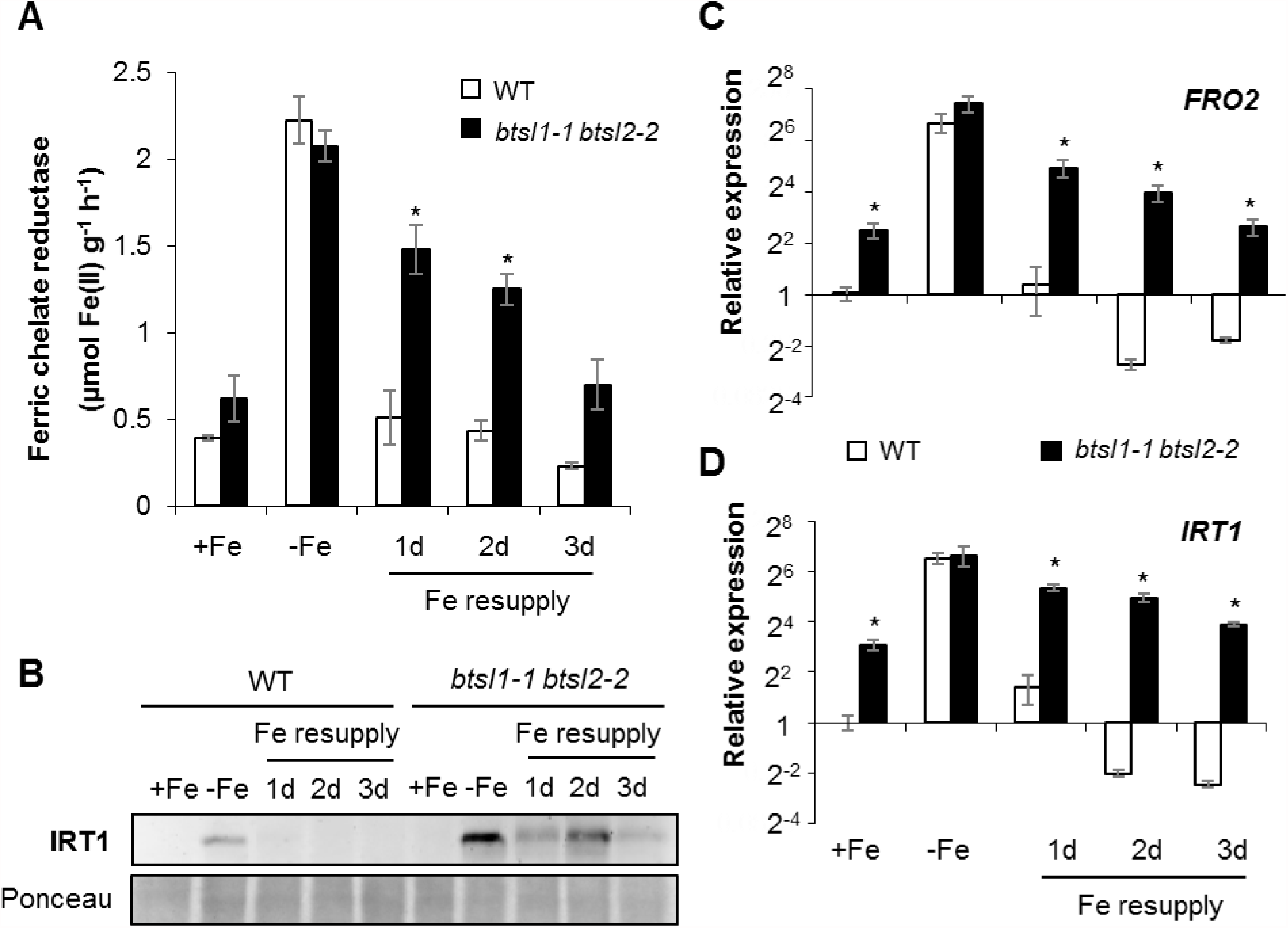
The *btsl* double mutant maintains FRO2 and IRT1 upon Fe resupply. **(A)** Ferric chelate reductase activity of wild-type and *btsl1-1 btsl2-2* mutants. Bars represent the mean of 3 biological replicates (n=5 seedlings/assay) ± SD (* p<0.05 using a two-tailed t-test). **(B)** Immunoblot of IRT1 protein levels in root samples in the Fe resupply treatment. **(C, D)** Expression of *FRO2* (C) and *IRT1* (D) in roots from wild-type and *btsl1-1 btsl2-2* seedlings determined by quantitative RT-PCR. All values are relative to wild type +Fe. Values are the mean of 3 biological replicates ± SE (* p<0.05 using a two-tailed t-test).

IRT1 protein levels in wild-type plants were strongly increased under Fe deficiency, and were nearly below detection after one day of Fe resupply. In the *btsl* double mutant, IRT 1 was not detected under standard conditions (+Fe), but after three days of Fe deficiency was present at much higher levels than in wild-type roots. Upon Fe resupply, the IRT1 protein levels fluctuated but remained high overall (Figure 6B). The sustained presence of FRO2 and IRT1 would explain the increased Fe accumulation in the *btsl* double mutant following the Fe-deficiency-resupply treatment.

To investigate which step of FRO2 and IRT1 expression is misregulated, we analysed transcript levels of *FRO2* and *IRT1* by RT-qPCR. Interestingly, under control conditions (+Fe), *FRO2* and *IRT1* transcripts levels were already increased 5 to 8-fold, respectively, in the *btsl* double mutant compared to wild type (Figure 6C, D), although no increase in enzymatic activity or protein levels was observed (Figure 6A, B). After three days of Fe deficiency, transcription of *FRO2* and *IRT1* was strongly upregulated by a similar magnitude in both mutant and in wild type. However, upon Fe resupply, *FRO2* and *IRT1* transcript levels remained high in the *btsl* double mutant, matching the sustained FRO2 activity and IRT 1 protein levels observed (Figure 6C). In contrast, in wild type, *FRO2* and *IRT1* transcripts were downregulated on day 2 and 3 after resupply compared to the initial levels (Figure 6C). Taken together, these data show that the transcriptional control of *FRO2* and *IRT1* is altered in the *btsl* double mutant and is disconnected from the Fe concentration in the medium.

### The transcriptional regulator FIT-bHLH38/39 is mis-regulated in the *btsl* double mutant

The transcription of *FRO2* and *IRT1* is directly regulated by FIT together with one of four partially redundant bHLH transcription factors (bHLH38, bHLH39, bHLH100, bHLH101). To investigate if FIT is misregulated, its protein and transcript levels were followed in wild-type and mutant plants subjected to the Fe deficiency-resupply treatment. Under control conditions, FIT was not detected by protein blot analysis in either wild-type or *btsl1-1 btsl2-2.* FIT protein levels were strongly induced after three days of Fe deficiency in wild-type seedlings, and even more so in *btsl* double mutant seedlings (~3-fold more than wild type, Figure 7A). Upon Fe resupply, FIT protein levels fell below the limit of detection in wild type, but were partially sustained in *btsl* roots, which could explain the increase in *FRO2* and *IRT1* transcripts (Figure 6C). In contrast, the transcriptional regulation of *FIT* was similar in wild-type and *btsl* seedlings (Figure 7B). This suggests than the protein levels of FIT may be directly or indirectly controlled by BTSL1 and/or BTSL2 under Fe deficiency, as well as upon Fe resupply.

**Figure 7.**
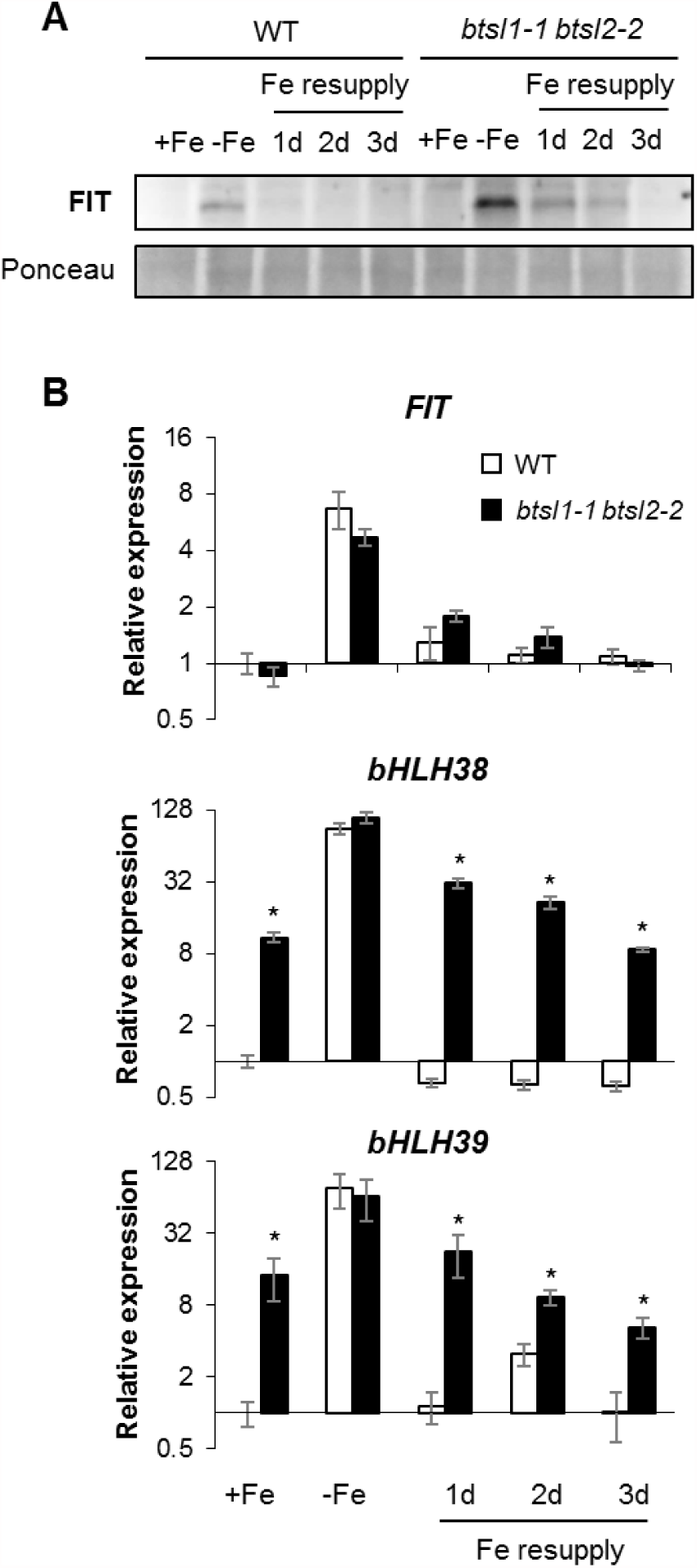
FIT protein and *bHLH38, bHLH39* transcript levels are misregulated in the *btsl* double mutant upon Fe resupply. (A) Immunoblot of FIT protein levels in root samples of the Fe deficiency-resupply treatment. (B) Expression of *FIT, bHLH38* and *bHLH39* in roots from wild-type and *btsl1-1 btsl2-2* seedlings determined by quantitative RT-PCR. All values are relative to wild type +Fe. Values are the mean of 3 biological replicates ± SE (* p<0.05 using a two-tailed t-test).

As already stated, FIT functions as a heterodimer, therefore we tested *bHLH38* and *bHLH39* transcript levels as a representative of all four redundant bHLH transcription factors from the subgroup Ib. Interestingly, under control conditions, *bHLH38/39* transcripts levels were increased 8-fold in the *btsl* double mutant compared to wild type (Figure 7B). After three days of Fe deficiency, the upregulation was similar in both genotypes, but upon Fe resupply, *bHLH38/39* transcript levels remained high in the *btsl* double mutant, matching the patterns observed for *FRO2* and *IRT1* transcripts (Figure 7B). In wild type plants, as soon as day 1 after resupply *bHLH38/39* transcripts were downregulated below initial levels (Figure 7C).

In summary, our data show that BTSL1 and 2 are required for FIT protein turnover, and for repressing transcription of *bHLH38/39.*

### BTSL1 and BTSL2 interact with FIT *in vitro*

Next, we tested if BTSL proteins can physically interact with FIT, a prerequisite for a direct target of ubiquitin-mediated protein degradation. A standard yeast 2-hybrid approach was not possible because of self-activation by the individual proteins. Therefore, we opted for an *in vitro* approach. The C-terminal CHY/RING Zn-finger domain of BTSL1 or BTSL2 was fused to MBP (maltose binding protein) for stabilization, and expressed in *E. coli.* FIT was also produced in *E. coli* with a Myc tag at the C-terminus. bHLH39:Myc and bHLH40:Myc were produced as controls. The protein interactions were studied by Far-Western blot analysis (Figure 8A) to overcome issues with the stability of the transcription factors. *E. coli* extracts with FIT, bHLH39 or bHLH40 were separated by SDS-PAGE and immobilized on nitrocellulose membrane. Immuno-detection of the Myc tag confirmed the presence and protein levels of FIT, bHLH39 and bHLH40 (Figure 8B and Supplementary Figure S6, left panels). Purified MBP, MBP:FIT or MBP:BTSL proteins were incubated with the membrane, followed by antibody detection of the MBP tag. MBP:FIT bound strongly to bHLH39 (35 kDa band) and weakly to itself (55 kDa band), as previously reported (Yuan et al 2008). The immuno-signals were clearly separated from an aspecific band in the MBP control. MBP:BTSL1c and MBP:BTSL1c bound to FIT, but not to bHLH39. A separate set of experiments showed that the BTSL proteins also did not interact with bHLH40 (Supplementary Figure S6), better known as the transcription factor INDEHISCENT involved in abscission but unrelated to iron homeostasis (Liljegren et al., 2004). These data suggest that both BTSL1 and BTSL2 bind FIT to target it for degradation.

**Figure 8.**
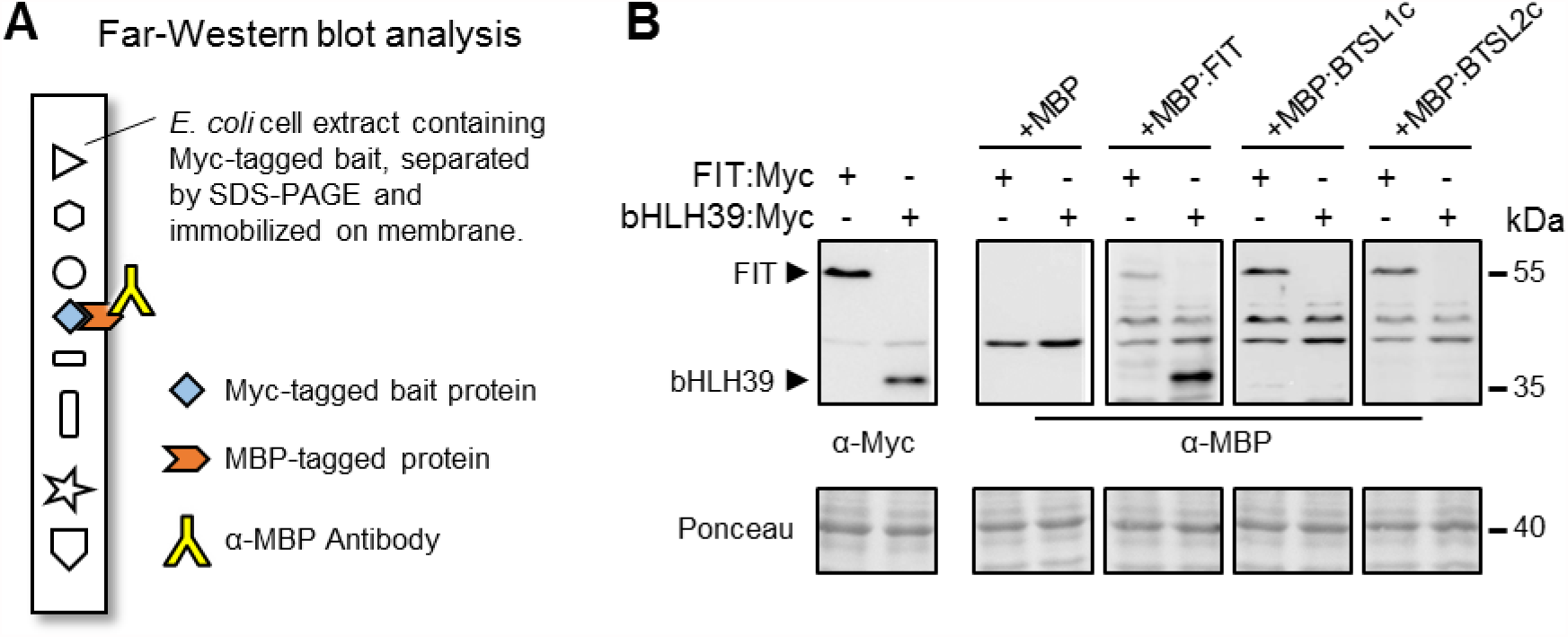
The CHY/RING Zn finger domain of BTSL proteins interact with FIT. (A) Diagram of Far-Western blot analysis. Bait proteins with a Myc-tag were produced in *E. coli* and protein extracts separated by SDS-PAGE gel followed by transfer to nitrocellulose membrane. Blots were incubated with purified MBP-tagged proteins of interest, then labelled with anti-MBP antibodies. (B) Far-Western blots analysis showing direct protein interactions between BTSL proteins and FIT. Bacterial protein extracts with FIT:Myc and bHLH39:Myc were immobilized on nitrocellulose, and the blots were incubated with recombinant MBP, MBP:FIT and the C-terminal domains of BTSL1 and BTSL2, fused to MBP (MBP:BTSL1c and MBP:BTSL2c). FIT is known to form a heterodimer with bHLH39. Ponceau staining shows equal protein loading.

## DISCUSSION

In this study, we show that BTSL1 and BTSL2 have a clearly distinct function from BTS, and are not simply redundant paralogs. *BTSL* genes have evolved specifically in dicotyledonous plants, they are root specific and co-regulated with the Fe uptake machinery. We show that the partially redundant BTSL proteins negatively control Fe uptake in the plant, which is exacerbated under Fe deficiency-resupply conditions. Under these conditions, BTS does not play any role. A closer look at the expression patterns of the *BTSL1, BTSL2* and *BTS* genes suggest a demarcation by the Casparian strip, an effective barrier against nutrient overload of plants. The two *BTSL* genes are predominantly expressed in the root epidermis and cortex, whereas BTS is expressed inside the root stele and in the shoot. *BTSL2* is also expressed in the stele in the differentiation zone, but this is where the Casparian strip is not yet formed or still permeable (Nasser et al., 2012; Barberon et al., 2016). Moreover, the *BTSL2* promoter-GUS expression was analysed under Fe deficient conditions, when suberinization of the Casparian strip is suppressed to maximize metal and nutrient uptake (Barberon et al., 2016). We propose a model where BTSL proteins act as a primary defense mechanism against excess Fe uptake in the root. A second defense barrier against Fe overload would be the BTS protein in the stele and leaf tissues, behind the Casparian strip (Figure 9).

**Figure 9.**
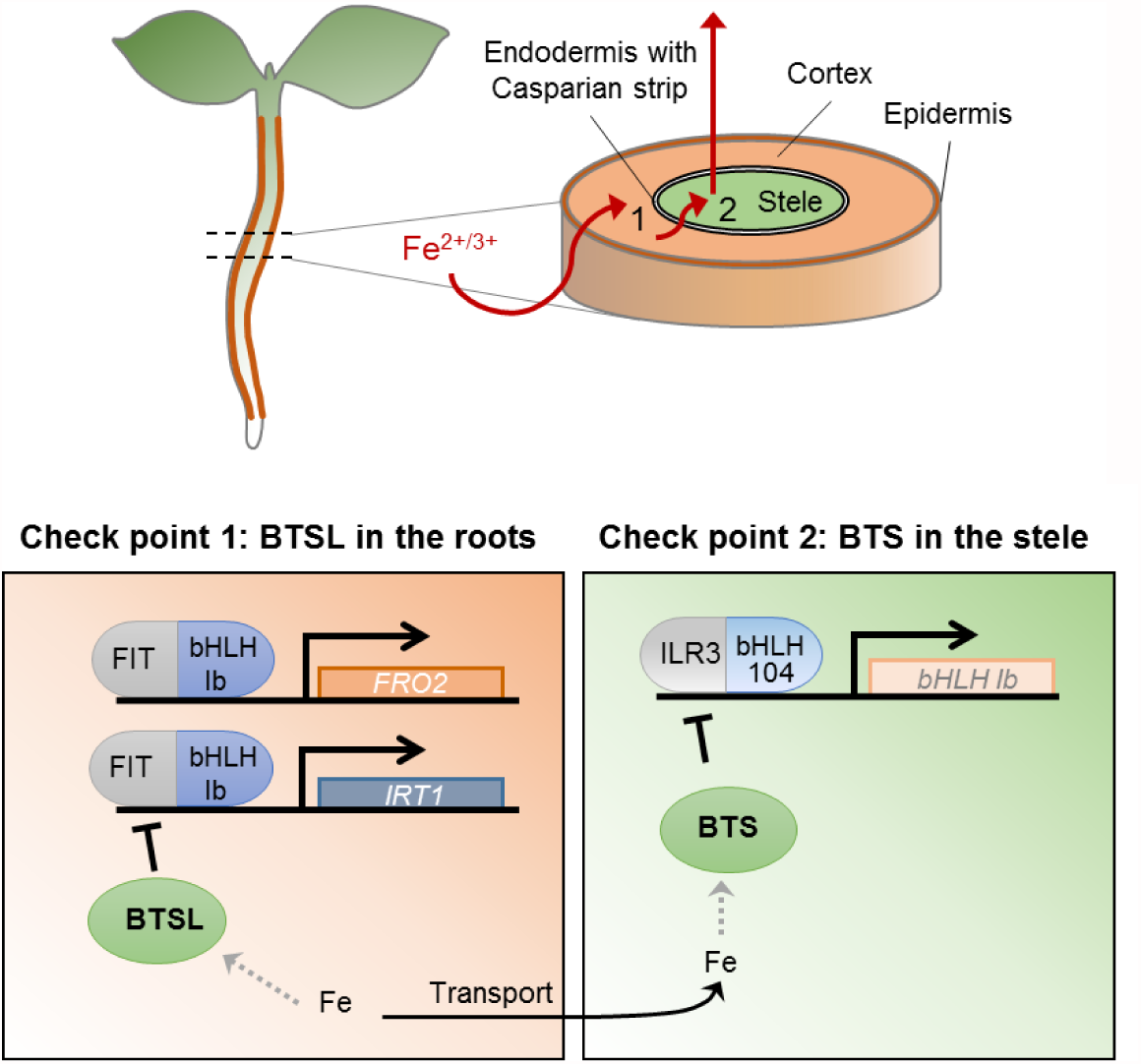
Proposed mode of action of BTSL and BTS in different parts of the plant. A model of how BTSL and BTS function to control Fe homeostasis. Fe levels are first sensed in the epidermal and cortex cells, where BTSL proteins respond by targeting FIT protein for degradation, thus controlling the uptake of Fe by FRO2 and IRT1 (and other proteins). A second checkpoint to protect plants from excess iron is present in the stele, stem and leaves, where BTS negatively controls ILR3 and bHLH104 protein levels (Selote et al., 2015). How iron affects the activity of BTS/L is not yet known (gray dashed arrows).

The different functions of BTSL1/2 and BTS suggest that the E3 ligases target different proteins for degradation. Their C-termini are rather unique among E3 ligases, consisting of a combination of CHY/RING-type Zn finger domains, as shown in NMR structures of the mammalian Pirh2/RCHY1 homolog (Sheng et al., 2008) which shares 45% amino acid identity with the C-termini of BTS/L. Pirh2/RCHY1 binds to the tetrameric form of the p53 transcription factor and targets it for degradation (Leng et al., 2003; Sheng et al., 2008). In Arabidopsis, 7 proteins share this conserved Pirh2-like domain: AT3G62970, AT5G18650, AT5G25560, AT5G22920; and BTSL1, BTSL2 and BTS. The best characterized is MYB30-INTERACTING E3 LIGASE 1 (MIEL1, AT5G18650), which targets the MYB30 transcription factor for degradation (Lee and Seo, 2016). Thus, it is likely that the unique CHY/RING-finger domain facilitates interaction with transcription factors for ubiquitination and degradation. BTS was found to interact and induce the degradation of the bHLH transcription factors ILR3 and bHLH115 (Selote et al., 2015), but direct ubiquitination of these two putative targets remains to be shown. It has recently been shown that HRZ1, the BTS homolog in rice, targets PRI1 (Positive Regulator of Iron homeostasis1) for degradation (Zhang et al., 2017). The bHLH transcription factor PRI1 is the homolog of Arabidopsis ILR3 and promotes the expression of *OsIRO2* and *OsIRO3,* which are orthologous to *bHLH38/39* and *PYE,* respectively (Kobayashi et al., 2013).

Our data suggest that the BTSL proteins target FIT for degradation. Firstly, the expression patterns of *FIT* and *BTSL* genes overlap in the epidermis and cortex (Figure 3; Colangelo and Guerinot, 2004; Jakoby et al., 2004). Secondly, FIT protein levels are increased in the *btsl* double mutant (Figure 7). Thirdly, the C-termini of BTSL1 and BTSL2 interact with FIT in vitro (Figure 8). It has previously been reported that FIT is regulated post-transcriptionally *via* proteasomal degradation (Sivitz et al., 2011), being turned over in response to ethylene (Lingam et al., 2011). Our data suggest that the BTSL proteins act as the E3 ligases responsible for ubiquitination of FIT which is then degraded by the 26S proteosome. At the moment, we cannot rule out that BTSL proteins may have other targets. The misregulation of *bHLH38* and *bHLH39* expression cannot be explained by lack of FIT protein degradation, because the transcriptional activitators of bHLH38/39 are ILR3 and bHLH104 (Zhang et al., 2015). Either BTSL also target ILR3 and/or bHLH104 for degradation specifically in those root tissues where BTS is not expressed, or mutation of the *BTSL* genes has an indirect effect on the BTS regulon. A comprehensive study of the direct targets of BTSL and BTS proteins will clarify these questions, although the experimental approaches for determining E3 ligases targets are not straight forward (Iconomou and Saunders, 2016).

Where the Fe status is sensed is still under debate (Kobayashi and Nishizawa, 2014; Gayomba et al., 2015). The observation that *BTSL* and *BTS* are expressed in different tissues support the hypothesis that Fe is sensed independently in different parts of the plant. Our data suggest that BTSLs act as local Fe sensors controlling Fe uptake. FRO2 activity and transcript levels are misregulated in the *btsl* double mutant after Fe resupply. The same happens with IRT1 protein and transcript levels, that remain high irrespective of the new Fe status. It is interesting to note the fluctuating levels of IRT1 protein after Fe resupply, which could be explained by its own internalization/degradation mechanism (Barberon et al., 2011; Shin et al., 2013). This protective mechanism seems to be disconnected from transcriptional regulation, with decreased levels of detectable IRT 1 protein one day after resupply. However, the sustained transcript levels then seem to overrule the post-translational control mechanism, and two days later detectable protein levels are almost as high as under Fe deficiency. On the inside of the Casparian strip, BTS could sense the Fe status in the shoot and control systemic signals that regulate Fe homeostasis. Both local and systemic controls of Fe homeostasis have been reported (Grusak and Pezeshgi, 1996; Schikora and Schmidt, 2001; Vert et al., 2003).

An open question still is of how the BTS/L proteins sense Fe and adjust their E3 ligase activity. The BTS/L proteins have conserved hemerythrin domains that are predicted to bind 2 Fe atoms per domain. Recombinantly expressed HRZ and BTS were found to bind 2 Fe and 1 - 5 Zn atoms (Kobayashi et al., 2013; Selote et al., 2015), which is far below the 6 Fe expected for 3 Hr domains and 9 Zn for the CHY/RING-type Zn finger domain. HRZ and BTS proteins seem to be rather unstable, and GFP fusions are not detected (Selote et al., 2015). Removal of the N-terminal domain with the three hemerythrin motifs led to a stable GFP fusion protein, suggesting that this domain regulates the stability of BTS. Based on *in vitro* data, Selote et al (2015) concluded that BTS protein is stabilized in the absence of Fe bound to the hemerythrin domains. However, this mechanism is the opposite of what is observed for FBXL5, where Fe binding to the Hr domain stabilized the protein (Salahudeen et al., 2009). The fact that the phenotypes in *btsl* mutants are particularly noticeable upon Fe resupply points towards a stabilization of BTSL protein by Fe, in a mechanism analogous to that of FBXL5. However, the changes in protein turnover in FIT and IRT1 under Fe deficiency suggest that BTSL proteins are also active under low Fe conditions. An in-depth biochemical characterization of the Hr domains is necessary to better understand the possible sensing mechanism of these proteins. Furthermore, the plant proteins have three hemerythrins as opposed to one in mammal FBXL5, suggesting cooperativity between the domains to fine tune the dynamic range of Fe binding affinities. BTSL and BTS may differ in this respect, given the different sequence of the second Hr domain (Supplementary Figure S2). Only when those questions are addressed will we be able to fully understand the Fe dependent regulation of the E3 ligase activity of BTS/BTSL/HRZ proteins.

## METHODS

### Co-expression analysis

The co-expression networks were produced using publicly available microarray data (NASCArrays; http://affymetrix.arabidopsis.info/) specific from roots and shoots using maccu software (http://maccu.sourceforge.net/). Networks were visualized using Cytoscape 3.0 (http://www.cytoscape.org/). BTSL1, BTSL2 and BTS were used as baits to retrieve coexpressed genes with a Pearson correlation coefficient >0.60 for BTSL1 and BTSL2, and >0.50 for BTS, and correlations between the retrieved genes were also calculated. Only those genes also present in the published ferrome (Schmidt and Buckhout, 2011) were retained.

### Phylogenetic analysis

Homologous sequences were found by using BLAST, with Arabidopsis BTS, BTSL1, BTSL2 and rice HRZ1 as queries in the Ensembl Plants database (http://plants.ensembl.org). Amino acid alignments were performed using Clustal Omega. A phylogenetic tree was plotted with BioNJ with the Jones-Taylor-Thornton matrix and rendered using TreeDyn 198.3. Species used: *Amborella trichocarpa, Arabidopsis thaliana, Brassica oleracea, Hordeum vulgare, Medicago truncatula, Oryza sativa, Physcomitrella patens, Populus trichocarpa, Solanum lycopersicum* and *Triticum aestivum.*

### Plant material

*Arabidopsis thaliana* ecotype Columbia (Col-0) plants were used as wild type. The T-DNA insertion lines were *btsl1-1* (SALK_015054); *btsl1-2* (SALK_032464); *btsl1-3* (SALK_117926); *btsl2-2* (SALK_048470); and *bts-1* (SALK_016526); from the Nottingham Arabidopsis Stock Centre. The T-DNA insertion sites were confirmed by sequencing. Double mutant lines were produced by crossing single insertion lines *btsl1-1* and *btsl1-3* with *btsl2-2.* Promoter sequences of *BTSL1* (-880 nt) and *BTSL2* (-2097 nt) were isolated by PCR from genomic DNA using primers BTSL1prom_GWF and BTSL1prom_GWR (BTSL1), and BTSL2pro_GWR and BTSL2pro_GWF (BTSL2). The promoter fragments were cloned upstream of eGFP-GUS in pBGWFS7 (Karimi et al., 2002) and the constructs were stably expressed in wild-type Arabidopsis. pXB2FS7 containing the CaMV 35S promoter upstream of eGFP-GUS was used as a positive control. The *btsl1-1 btsl2-2* double mutant line was complemented with tagged versions of BTSL1 and BTSL2. Full coding sequences of *BTSL1* and *BTSL2* were cloned using the primer pairs BTSL1_B1-F and BTSL1_B1-R, and BTSL2_B1-F and BTSL2_B1-R, respectively. These sequences were cloned upstream eYFP and eGFP, respectively, in the vectors pEarleyGate101 and pEarleyGate103 (Earley et al., 2006). All the sequences of the primers used are in Supplementary Table S3.

### Plant growth conditions

For Fe treatments, plants were grown on half Hoagland solution solidified with 0.8% (w/v) agar. The medium was composed of (in mM): K (3.5), Mg (1), Ca (2.5), NO_3_ (7.5), PO_4_ (1), and SO_4_ (1); and (in μM): Mn (9.14), B (46.3), Mo (0.12), Zn (2.4) and Cu (0.37). The pH was adjusted to 5.5 with 1mM 2-(N-morpholino)ethanesulfonic acid (MES). For Fe treatments Fe was added at the desired concentrations using FeEDTA. For Fe deficiency media, no Fe was added and medium was supplemented with 100 μM of 3-(2-pyridyl)-5,6-diphenyl-1,2,4-triazine sulfonate (ferrozine). For seed propagation, plants were grown on compost under long-day conditions, 16 h light at 22°C, 8 h dark at 20°C, with 80% humidity and a light intensity of 140 - 200 μmol m^-2^ s^-1^.

### Promoter GUS studies

Glucuronidase activity was detected as described (Jefferson et al., 1987), keeping the reaction time constant for equivalent samples. Seedlings were cleared in 100% ethanol, or fixed with a solution of 3.7% (w/v) formaldehyde, 5% (v/v) acetic acid, 50% (v/v) ethanol, followed by embedding in TechnoVit (Heraeus Kulzer GmbH, Germany), per manufacturer instructions. Transverse root sections (10 μm) were imaged on a LEICA DM6000 light microscope. The objectives used were x20/0.7 air or x40/0.85 air and images were collected with a LEICA DFC420C colour camera and processed with LEICA LAS-AF software.

### Plant phenotyping

Arabidopsis leaf pigments were extracted following a modified protocol from Abadía and Abadía (1993). Briefly, leaf pigments were extracted on ice with 80% acetone, and extracts were filtered and centrifuged to eliminate any particulate. Absorbance was measured at 470, 645, 662, and 750 nm wavelengths in a multiplate reader (ClarioStar; BMG Labtech, Germany). Mean values and standard deviations (SD) were obtained from three biological replicates.

Iron content in Arabidopsis leaves was determined by the Ferene method. Dried and powdered samples (4–8 mg) were mineralized at room temperature in 150 μL nitric acid (65%) for 2 days. One hundred and fifty microliters of H_2_O_2_ (30%) were added, and the solution was incubated 2 more days. The volume was adjusted to 500 μL with sterile water. A standard curve with Fe from 0 to 10 μg of Fe from ferric ammonium citrate was subjected to the same mineralization process. One hundred microliters of the mineralized samples were buffered with 1 mL of 15% ammonium acetate and reduced with 100 μL of 4% (w/v) ascorbic acid. Then, Fe(II) was chelated by adding 100 μL of 1.5% w/v 3-(2-pyridyl)-5,6-bis(5-sulfo-2-furyl)- 1,2,4-triazine (Ferene, SIGMA). Absorbance at 593 nm was measured in triplicates in a multiplate reader (ClarioStar, BMG Labtech).

To localize Fe in tissues, plants always grown on Fe-sufficient media or following a resupply after Fe deficiency induction were incubated with Perls’ stain solution (equal volumes of 2% [v/v] HCl and 2% [w/v] K-ferrocyanide) for 30 min. Seedlings were then washed three times with water, chlorophyll partially removed by incubation with 50% ethanol and then imaged with a Leica DM6000 microscope.

Fe chelate reductase activity was measured following Yi and Guerinot, (1996). Briefly, roots from 5 plants were blotted dried, weighed and incubated for 30 minutes in the dark with measuring solution containing 0.1 mM Fe(III)-EDTA and 0.3 mM ferrozine in distilled water. Fe^2+^ production was determined with a spectrophotometer measuring absorbance at 562 nm.

### Quantitative reverse-transcription PCR

Total RNA was extracted using the Plant RNeasy kit (Qiagen), followed by DNase treatment (Turbo DNase kit, Agilent). The integrity of RNA in all samples was verified using agarose gels, and RNA sample purity was analysed by comparing 260/230 nm and 260/280 nm absorbance ratios (Nanodrop 2000, Thermo Fisher). Only samples which passed these quality control checks were used. RNA was quantified using a Qubit 2.0 fluorometer (Thermo Fisher). RNA (4 μg) was reverse transcribed to cDNA using Superscript III (Thermo Fisher). RT-qPCR reactions were made using SensiFAST master-mix (Bioline), in 20 μl volumes, each with 20 ng of cDNA. Reactions were measured in a Bio-Rad CFX-96 real-time PCR system and cycled as per the Bioline protocol. Data were analysed using the Bio-Rad CFX Manager 3.1 software, and were normalized using primer efficiency. All data points are from three or more independent biological replicates, measured in three technical replicates (n=9). The house-keeping genes SAND (*AT2G28390*) and TIP41-like (*AT4G34270*) were used as reference genes, as they are unaffected by Fe levels in Arabidopsis (Han et al., 2013).

### Protein expression and immunoblot analysis

Roots from ten plants were ground in liquid nitrogen and suspended in acetone containing 10% (w/v) TCA, mixed thoroughly and precipitated for 2 hours at -20°C. Proteins were pelleted and washed three times with cold acetone and then dried prior solubilisation with a buffer containing 8 M urea, 0.5% SDS, 50 mM DTT and 1 mM PMSF. Proteins were separated on 12.5% SDS-PAGE gels, blotted into Nitrocellulose membranes and immunodetected using anti-Ferritin, anti-FIT and anti-IRT1 (Fan et al., 2014) antibodies and ECL Plus chemiluminescence substrate. Anti-Ferritin antibody was produced using purified pea ferritin for rabbit immunization. For anti-FIT antibody production, the coding sequence of FIT was fused to MBP, purified from *E. coli* and used for rabbit immunization.

### Recombinant protein production

For recombinant protein production, full coding sequences of *FIT* (*AT2G28160*), *bHLH39* (*AT3G56980*) and *bHLH40* (*AT4G00120*) were cloned from cDNA from Fe deficient Arabidopsis roots or genomic DNA, with a N-terminal His-tag and C-terminal Myc tag. Coding sequences were then inserted into the pET-15b vector using NdeI and BamHI restriction sites. The C-terminal coding regions of *BTSL1* (amino acids 933-1259) and *BTSL2* (amino acids 787-1254) were amplified from cDNA from Fe deficient Arabidopsis roots and cloned into the expression vector pMAL-c5X (NEB) using NotI-SalI and NcoI-SalI restriction sites, respectively. This generated an N-terminal fusion with maltose binding protein (MBP) to obtain MBP:BTSL1c and MBP:BTSL2c. A construct for MBP:FIT was produced in the same way. Plasmids were transformed into *Escherichia coli* Rosetta (NEB). Cells were grown in LB medium and protein expression induced with IPTG (Isopropyl β-D-1-thiogalactopyranoside) following manufacturer’s instructions for 4 h at 37°C. The recombinant MBP fusion proteins were purified using amylose resin. All primers used are listed in Supplementary Table S3.

### Far-western blot analysis

Soluble protein extracts of bacteria expressing FIT:Myc and bHLH40:Myc proteins were separated in a 10% SDS PAGE gel and blotted into a nitrocellulose membrane. Membranes were then blocked and incubated with purified MBP, MBP:FIT, MBP:BTSL1c or MBP:BTSL2c at 0.5 μM in TBS-tween buffer with 5% dry milk as blocking agent. After washing, proteins bound to the membrane were detected with anti-Myc (ab18185, Abcam) and ant-MBP (ab23903, Abcam) antibodies, respectively.

### Accession numbers

BTSL1 (AT1G74770); BTSL2 (AT1G18910); BTS (ATG3G18290);

OsHRZ2 (LOC_Os05g47780); OsHRZ1 (OS01T0689451_02);

TaHRZ1-A (TRIAE_CS42_3AL_TGACv1_195251_AA0646970);

TaHRZ1-B (TRIAE_CS42_3B_TGACv1_220625_AA0712080);

TaHRZ1-D (TRIAE_CS42_3DL_TGACv1_249432_AA0848520);

TaHRZ2-A (TRIAE_CS42_1AL_TGACv1_000799_AA0019360);

TaHRZ2-B (TRIAE_CS42_1BL_TGACv1_031945_AA0123190);

TaHRZ2-D (TRIAE_CS42_1DL_TGACv1_061110_AA0185650).

FIT / bHLH29 (AT2G28160); bHLH38 (AT3G56970); bHLH39 (AT3T56980); bHLH40 (AT4G00120)

## AUTHOR CONTRIBUTIONS AND ACKNOWLEDGMENTS

We are grateful for funding from BBSRC (BB/N001079/1) and the CEPAMS fund. J.R-C. was funded by a Marie Sklodowska Curie fellowship form the EU H2020 programme. We would like to thank Huilan Wu for help with Western blot analysis of FIT and IRT1; Benjamin Planterose for assistance with molecular cloning; the Bioimaging platform and Elaine Barclay for sectioning;

J.R.C. and J.B. conceived the project, performed experiments, analysed the data and wrote the paper. J.M.C., R.T.G. and I.K. performed experiments and analysed data. Y.C. and H.Q.L. contributed new analytical tools.

## Supplementary data

**Supplementary Figure S1.**
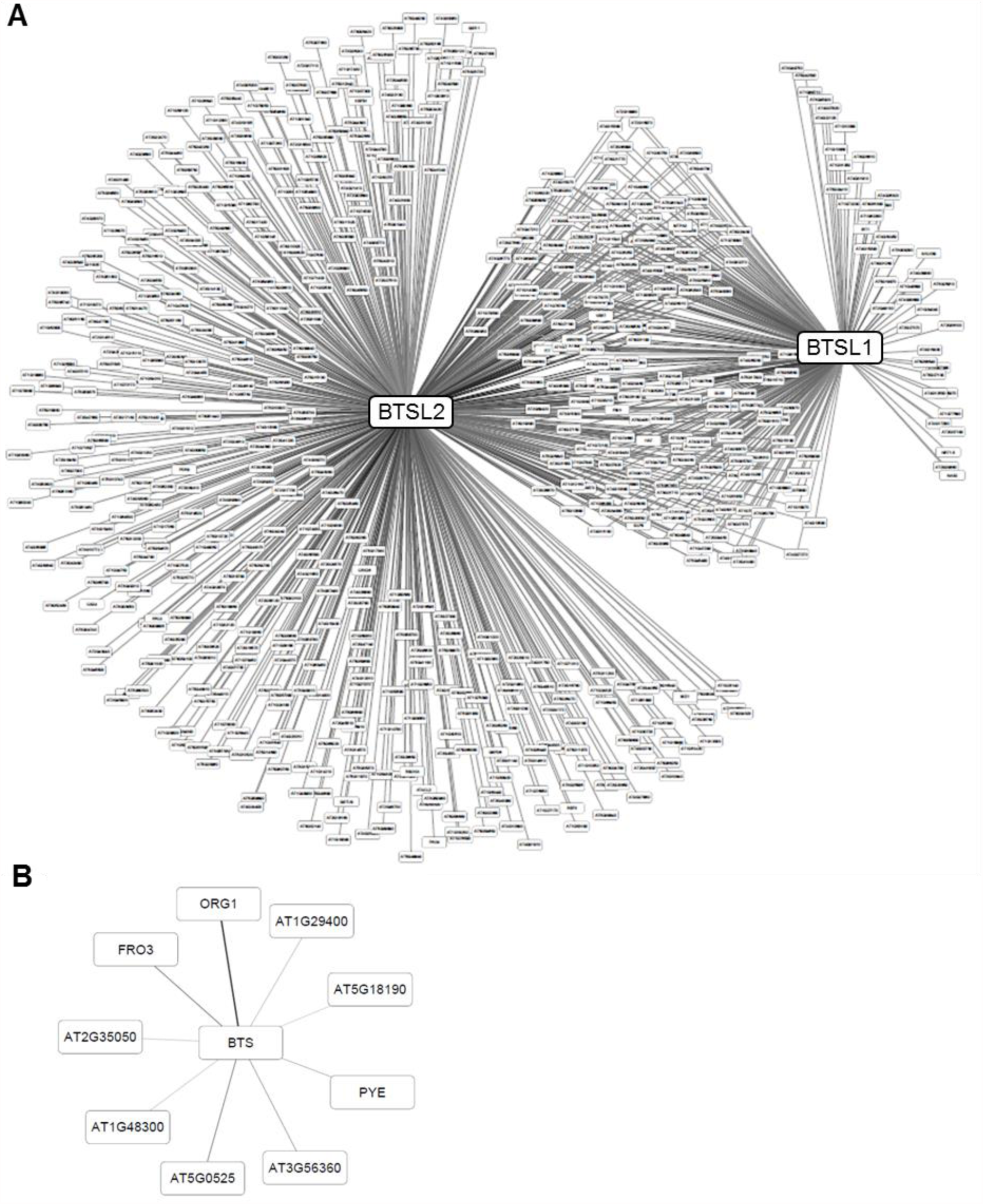
Full co-expression networks of *BTSL1, BTSL2* and *BTS*. **(A, B)** Co-expression analysis of genes that are co-regulated with (A) BTSL1 and BTSL2 in roots, and (B) BTS in shoots. The edge thickness relate to the Pearson’s correlation coefficient. See Supplemental Table S1 for descriptions of all genes and numerical values.

**Supplementary Figure S2.**
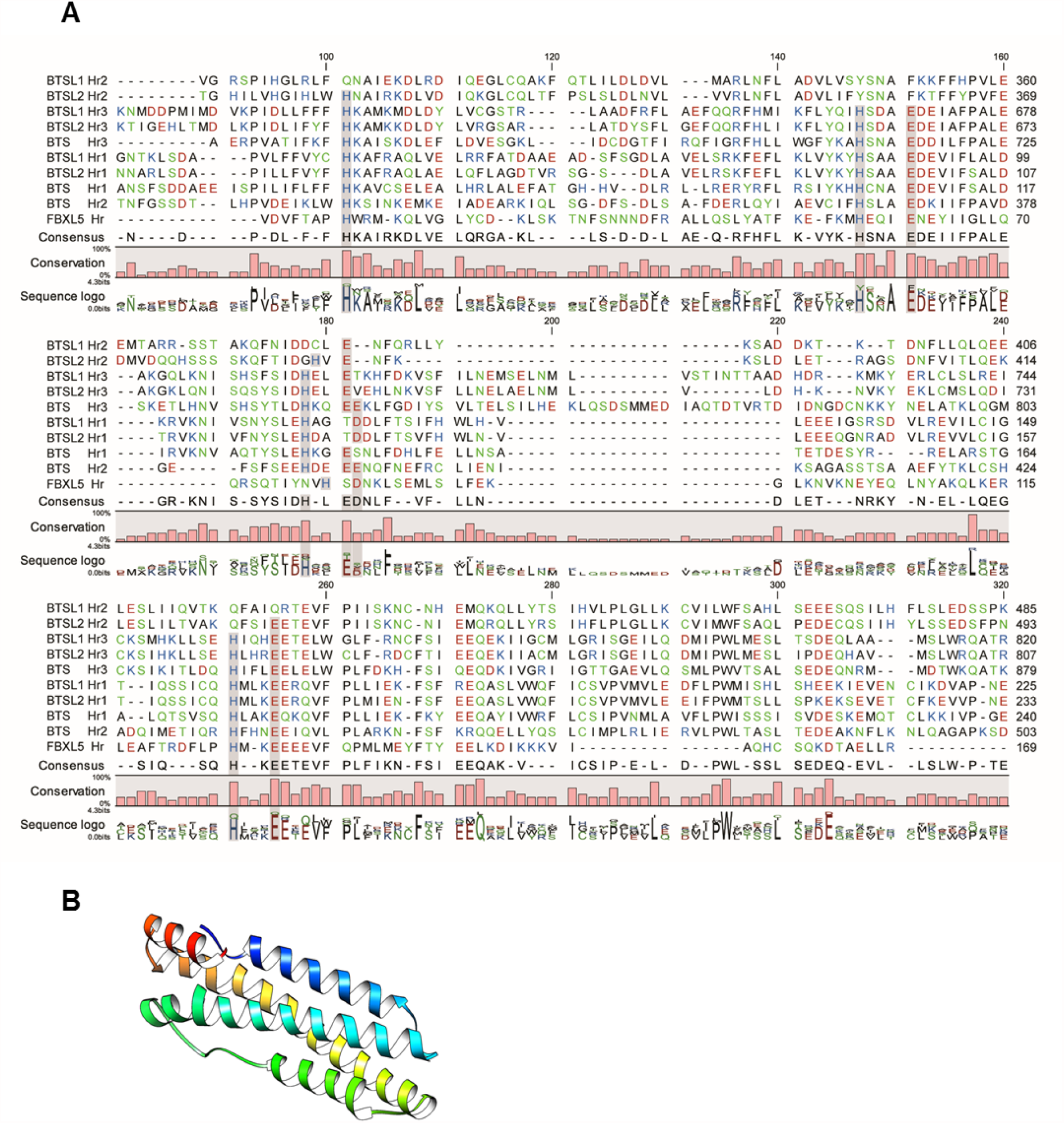
Alignment of hemerythrin domains and structure prediction for the second Hr domain of BTSL1. **(A)** Alignment of hemerythrin domains ofFBXL5, BTS, BTSL1 and BTSL2. **(B)** Predicted structure for the second Hr domain of BTSL1 using Phyre2 server.

**Supplementary Figure S3.**
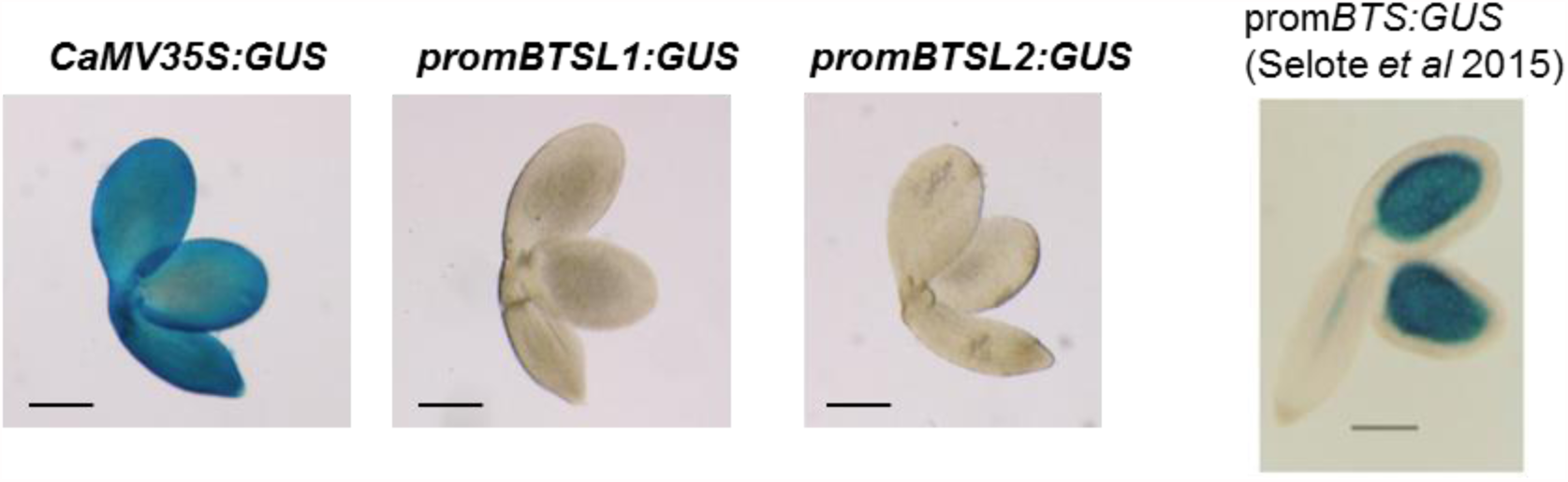
GUS staining of embryos. GUS activity staining of embryos from constitutive CaMV35S:GUS, promBTSLI: GUS and promBTSL2:GUS. Data for promBTS:GUS for comparison from Selote et al., 2015. Scale bar 200 μm.

**Supplementary Fig S4.**
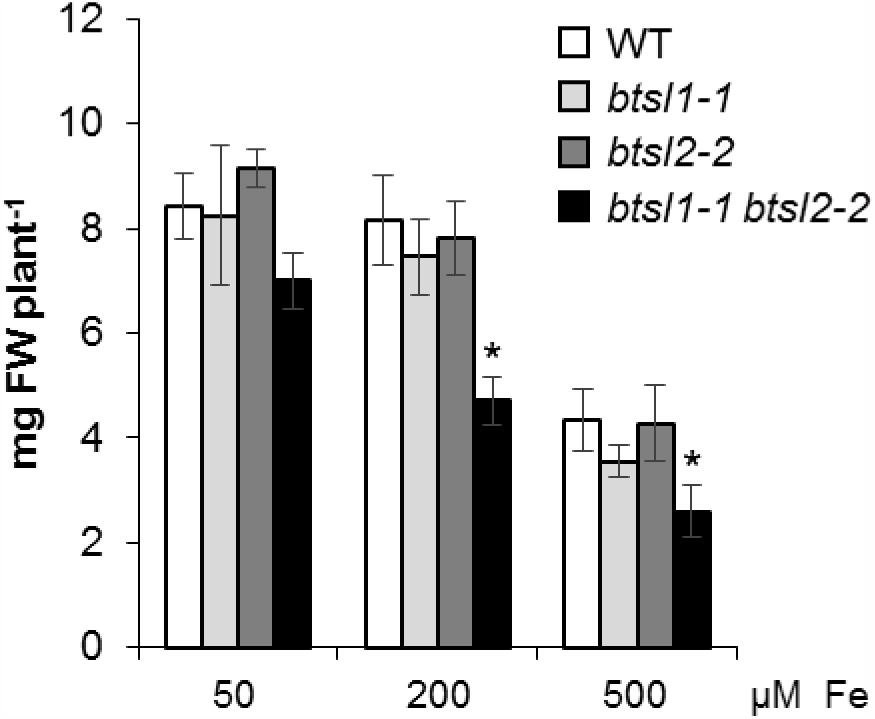
Fresh weight of WT and *btsl* mutant lines under Fe sufficiency and toxicity conditions. Shoot fresh weight of shoots from wild type (WT) and *btsl* mutant alleles, grown under control and Fe toxicity conditions as in Fig 4D. The bars represent the mean of n = 3 biol. reps x 10 seedlings ± SD. (* p<0.05 using a two-tailed *t*-test).

**Supplementary Fig S5.**
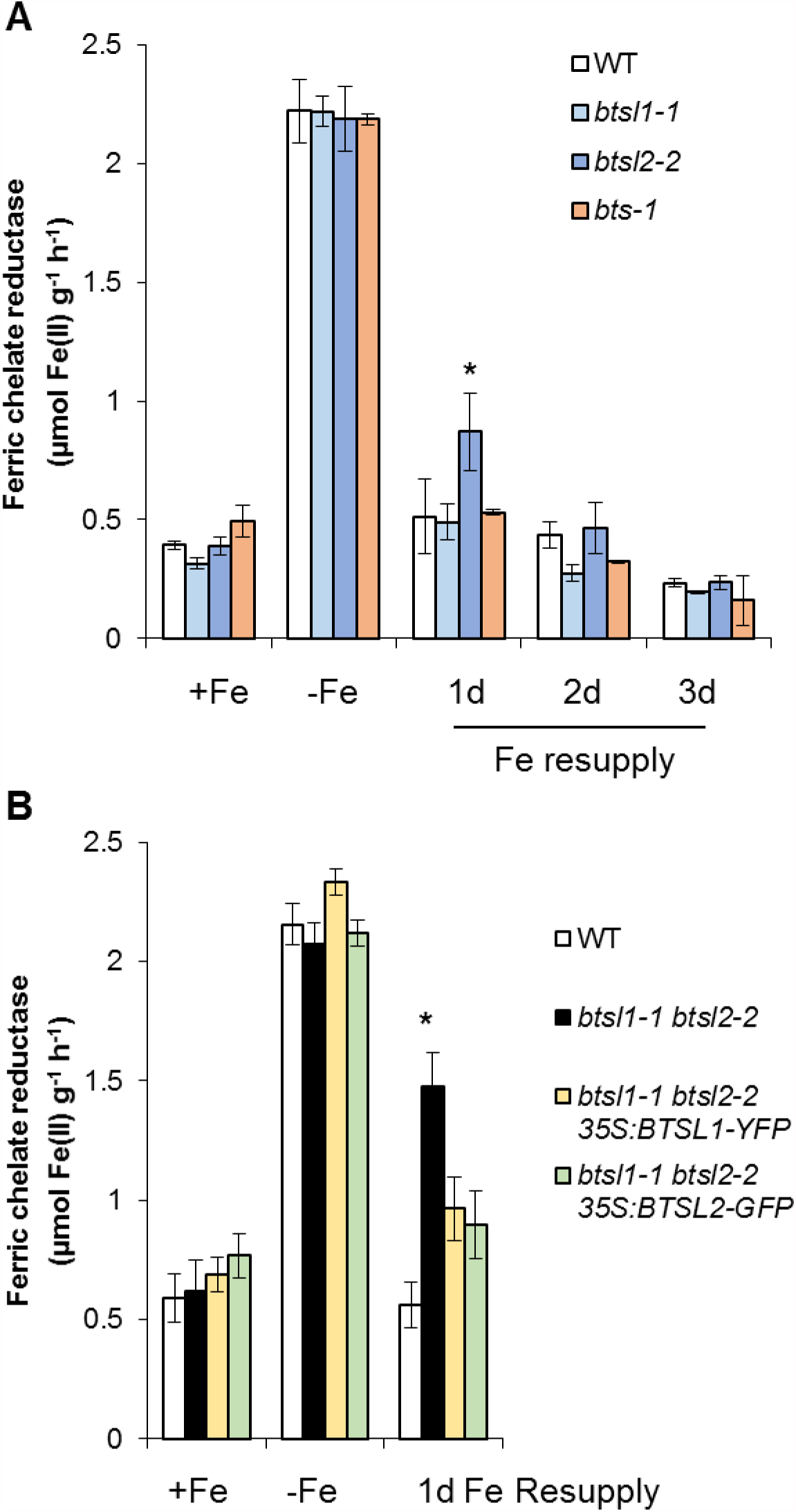
Fe reductase activity of *btsH-1, btsl2-2, bts-1* and complementation lines. (A) Ferric chelate reductase activity of *btsl 1-1, btsl2-2* and *bts-1* single insertion mutants under the Fe-defiency and resupply treatment. (B) Complementation of mutant alleles based on ferric chelate reductase activity. Wild type, *btsl1-1 btsl2-2*double mutant and *btsl1-1 btsl2-2* lines expressing 35S:BTSL1-YFP or 35S:BTSL2-GFP were grown under Fe sufficiency, Fe deficiency and resupplied with Fe for 1 day. The results are the mean value of 3 biological replicates +/- SE (*, p<0.05 using a two-tailed Student *t*-test).

**Supplementary Fig S6.**
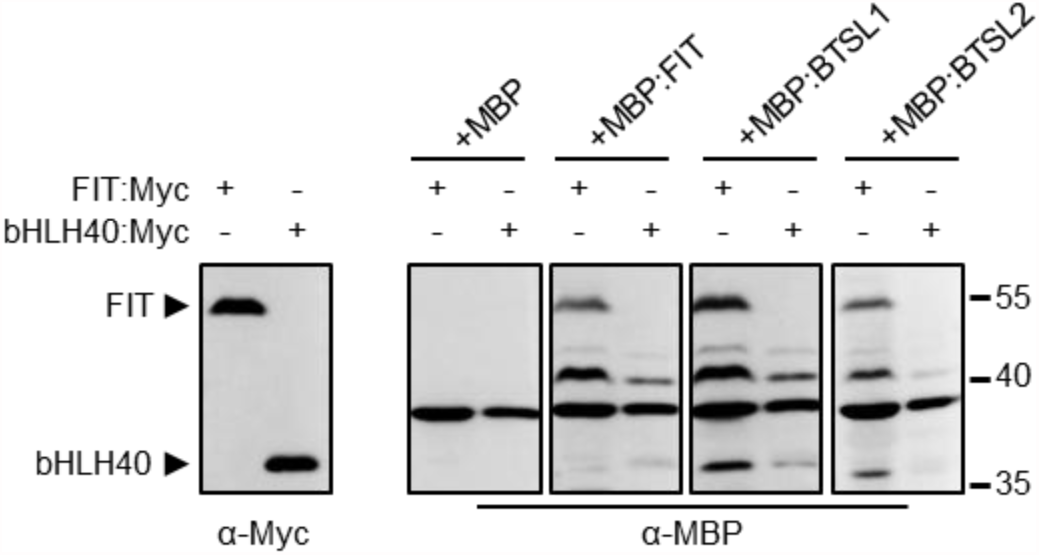
The CHY/RING Zn finger domain of BTSL proteins do not interact with bHLH40. Far-Western blots analysis showing no protein interactions between BTSL proteins and bHLH40. Bacterial protein extracts with FIT:Myc and bHLH40:Myc were immobilized on nitrocellulose, and the blots were incubated with recombinant MBP, MBP:FIT and the exterminai domains of BTSL1 and BTSL2, fused to MBP (MBP:BTSL1c and MBP:BTSL2c). FIT is known to not form a heterodimer with bHLH40.

**Supplementary Table S1.**
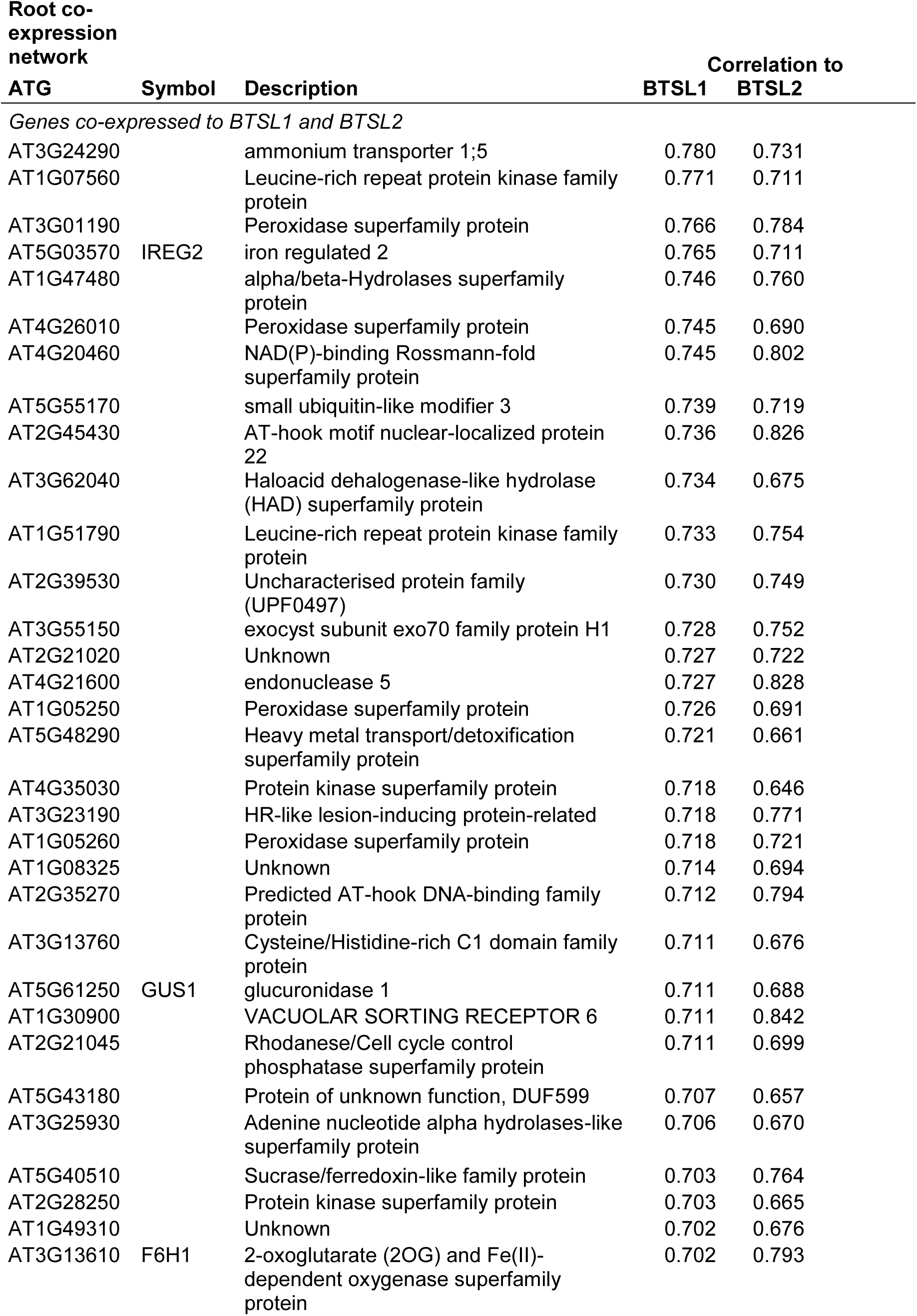
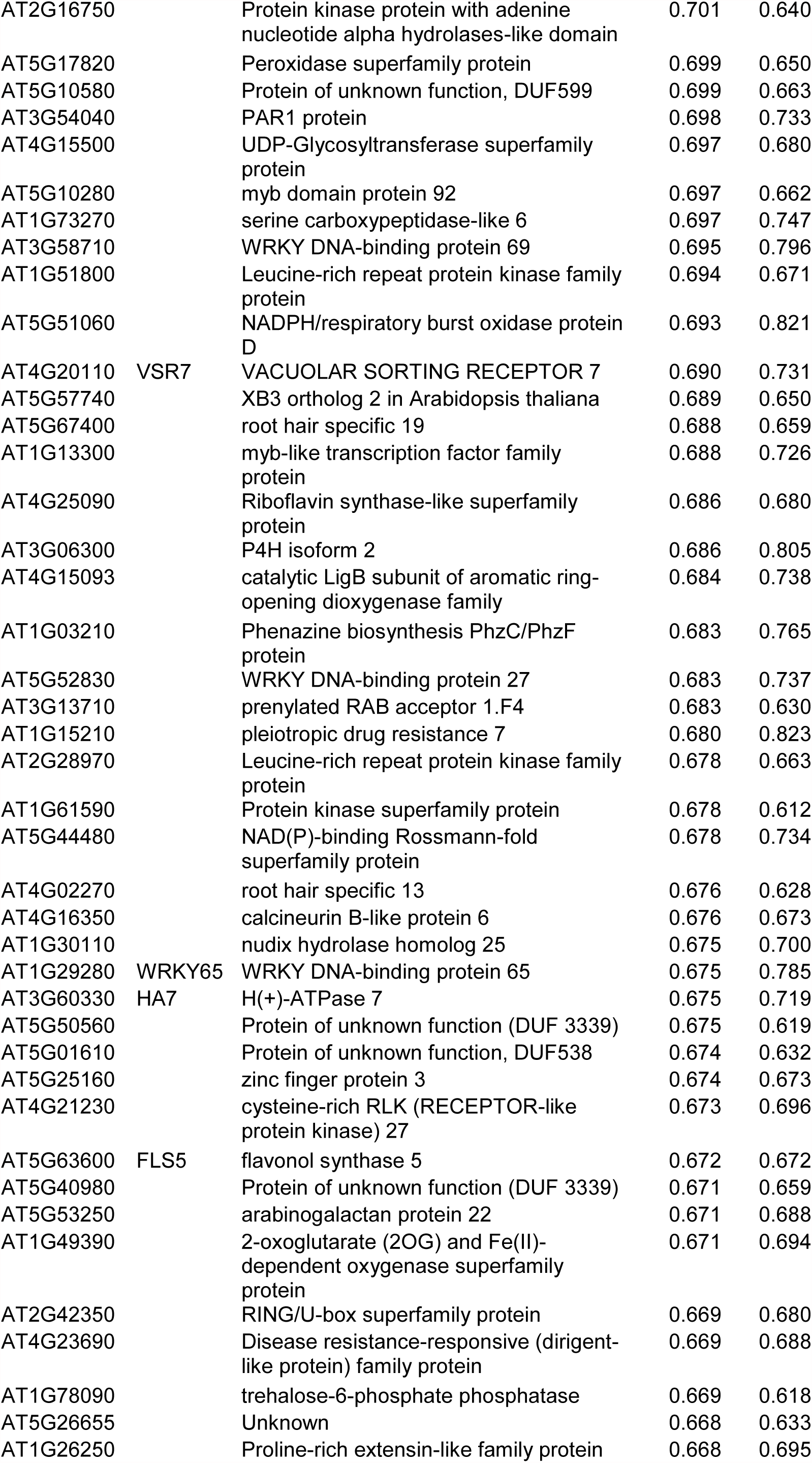
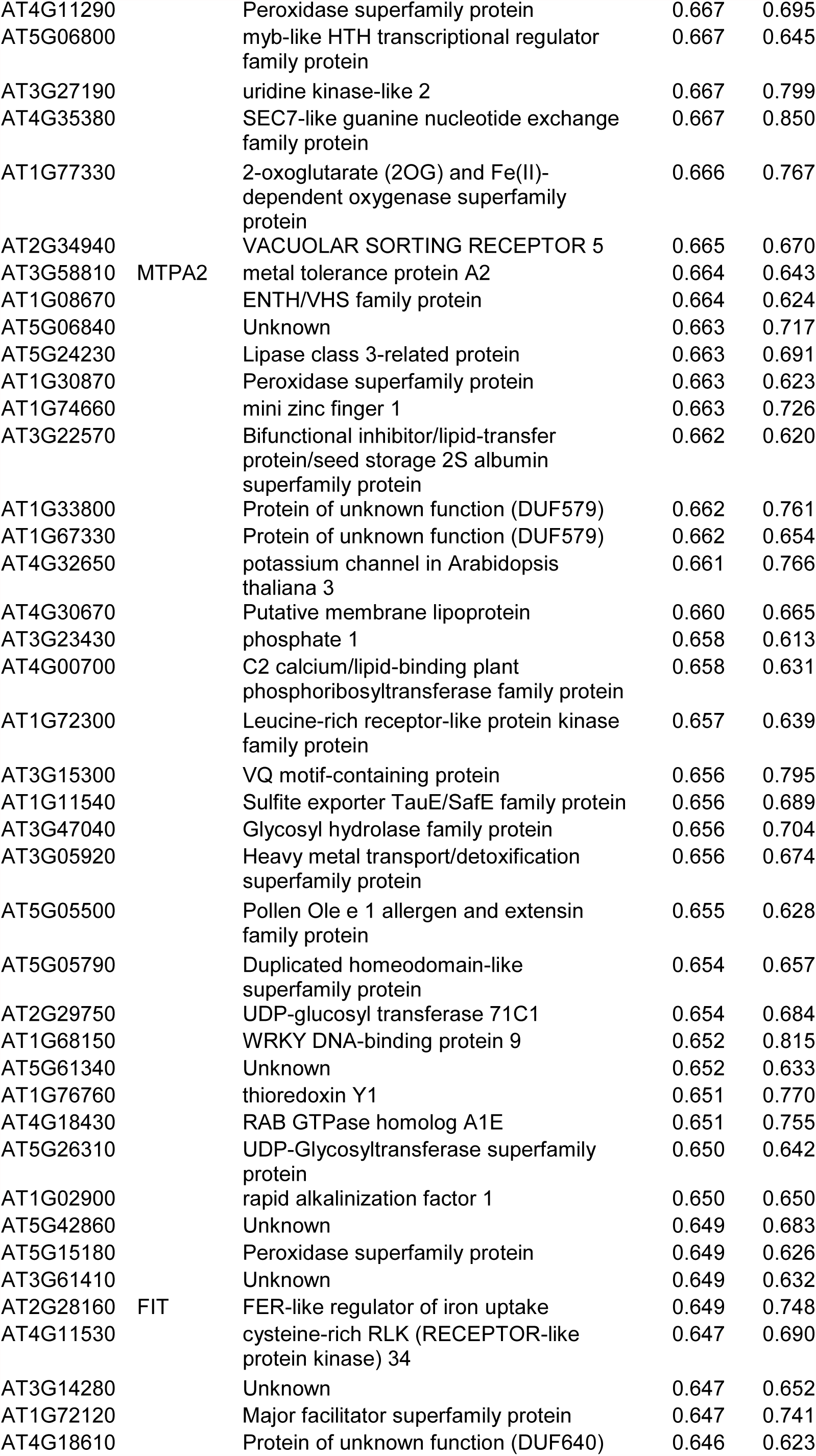
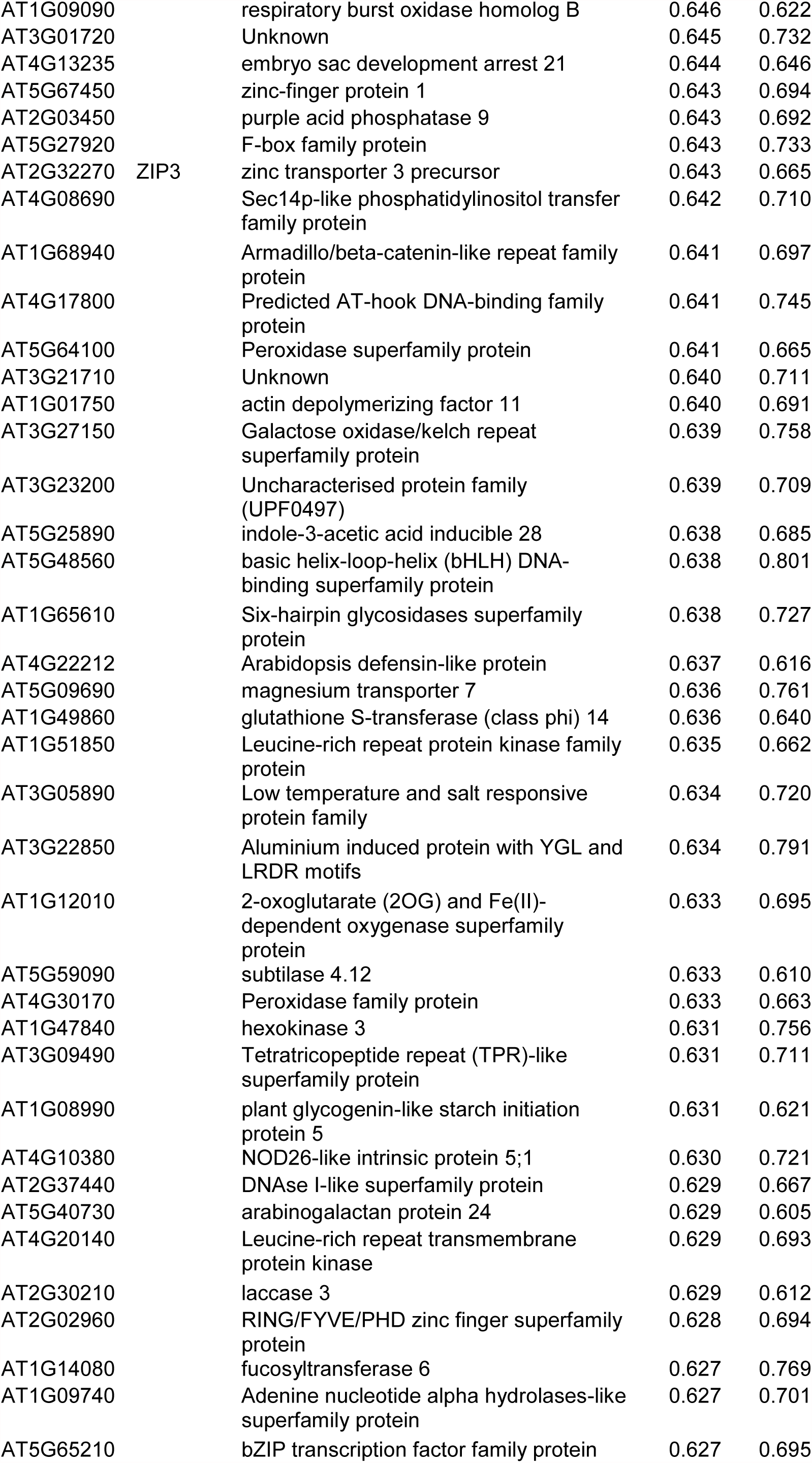
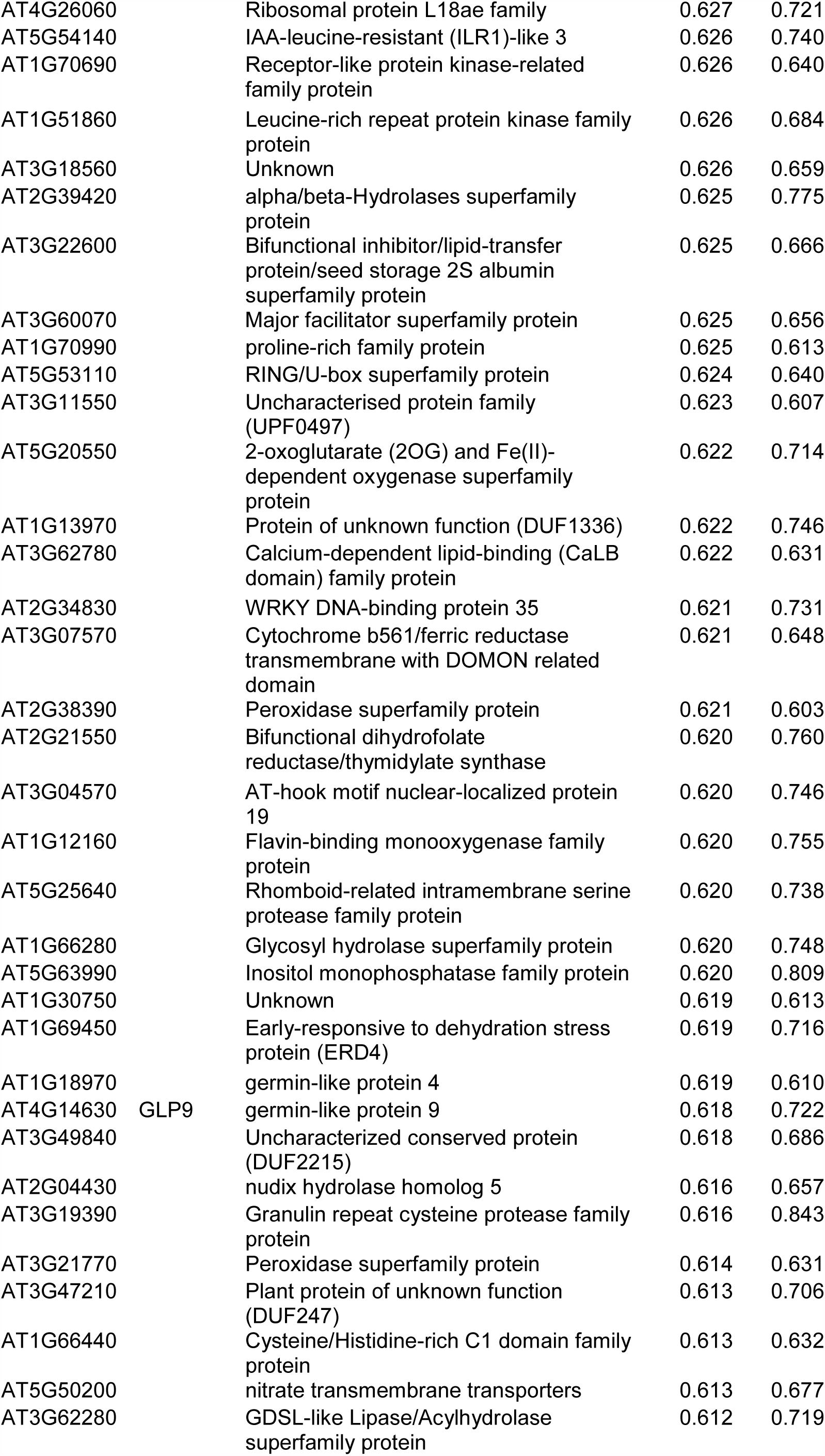
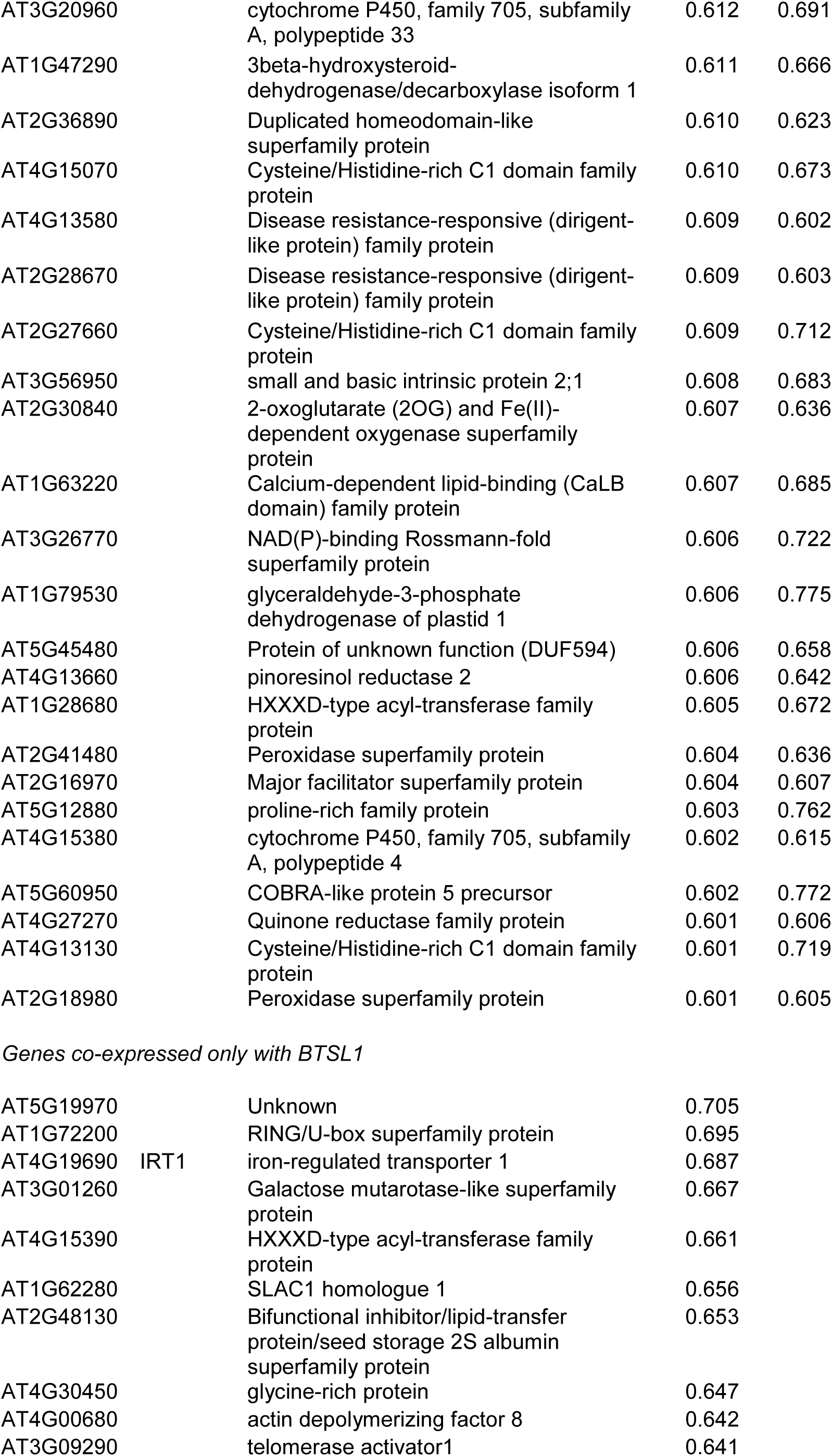
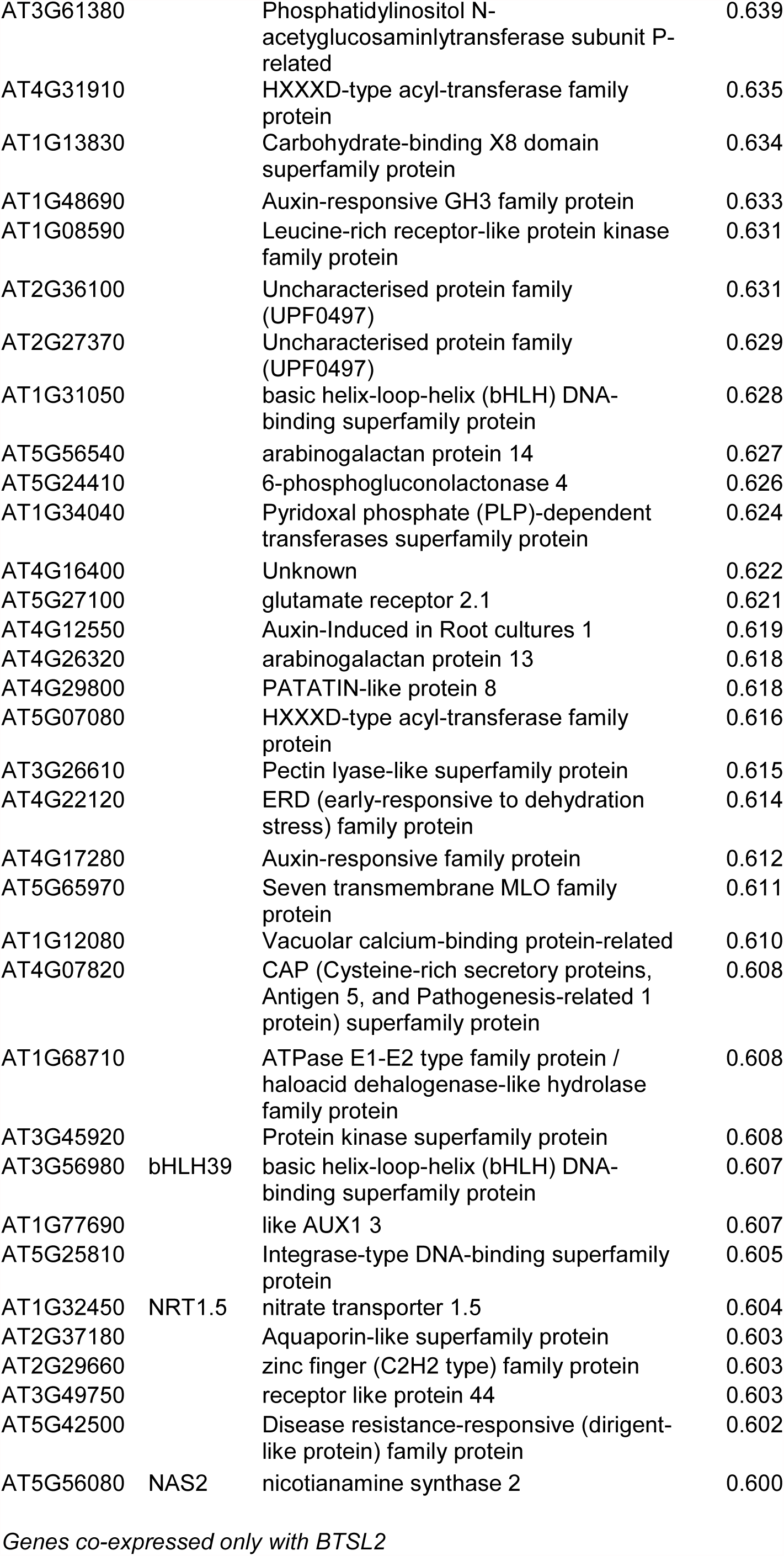
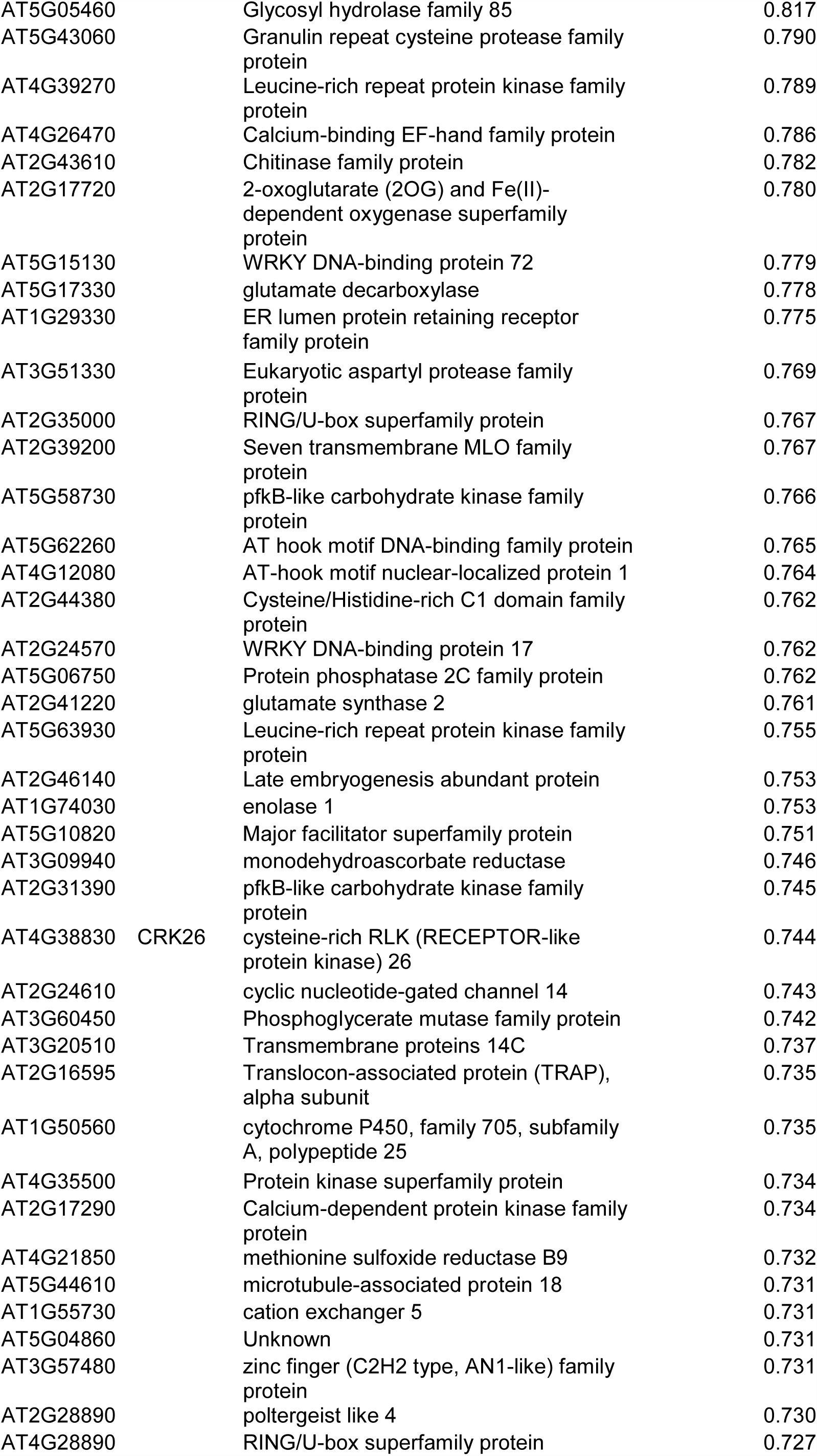
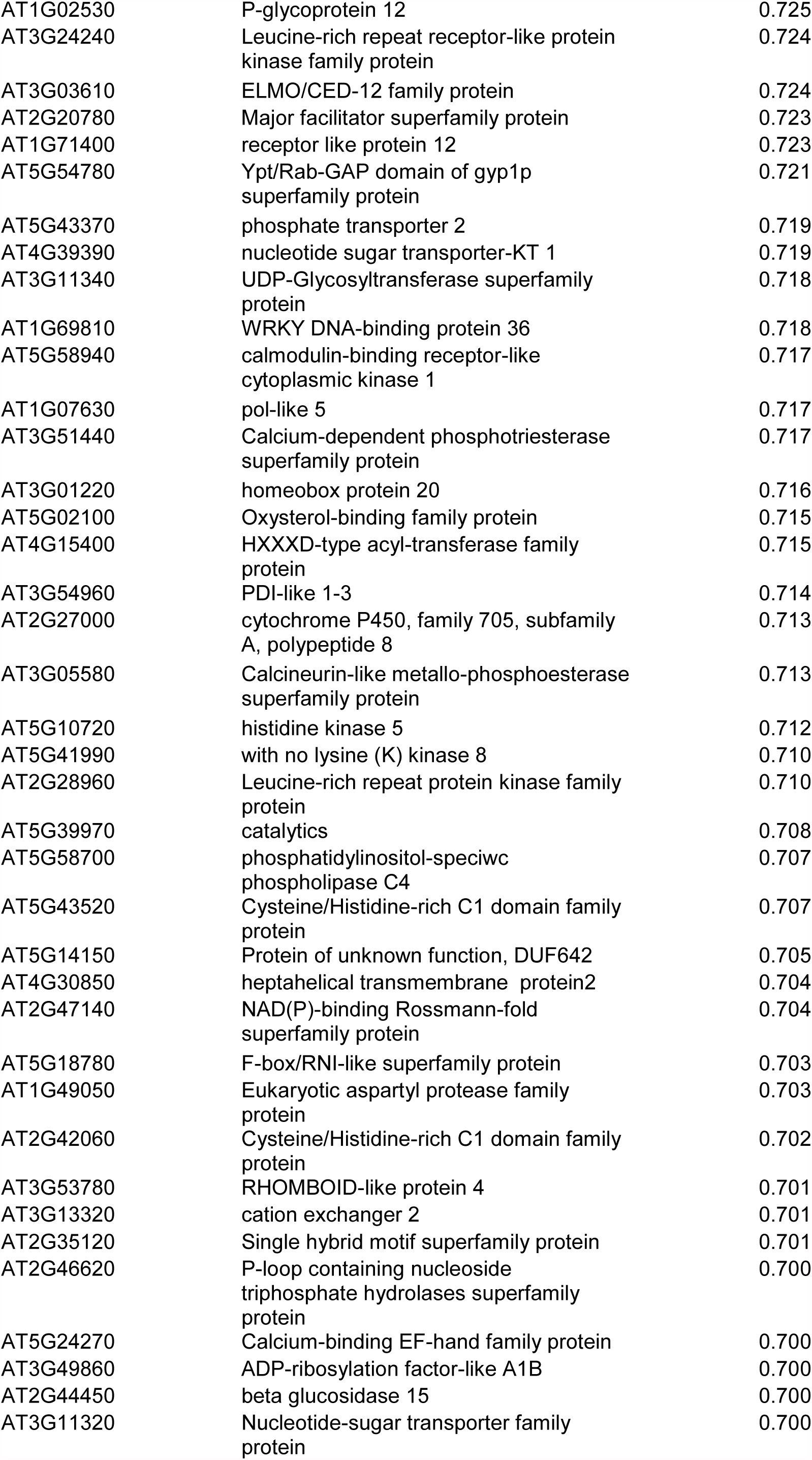
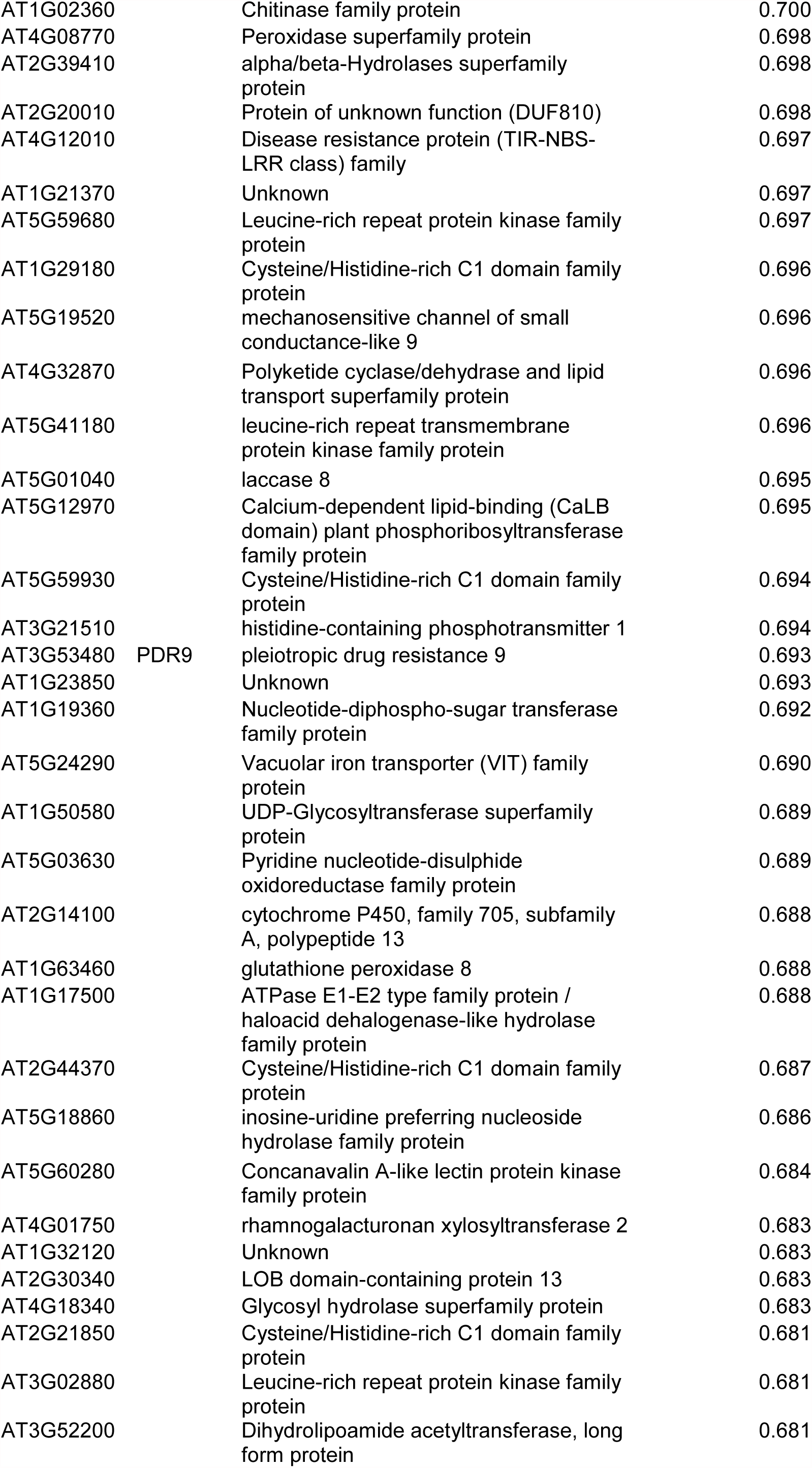
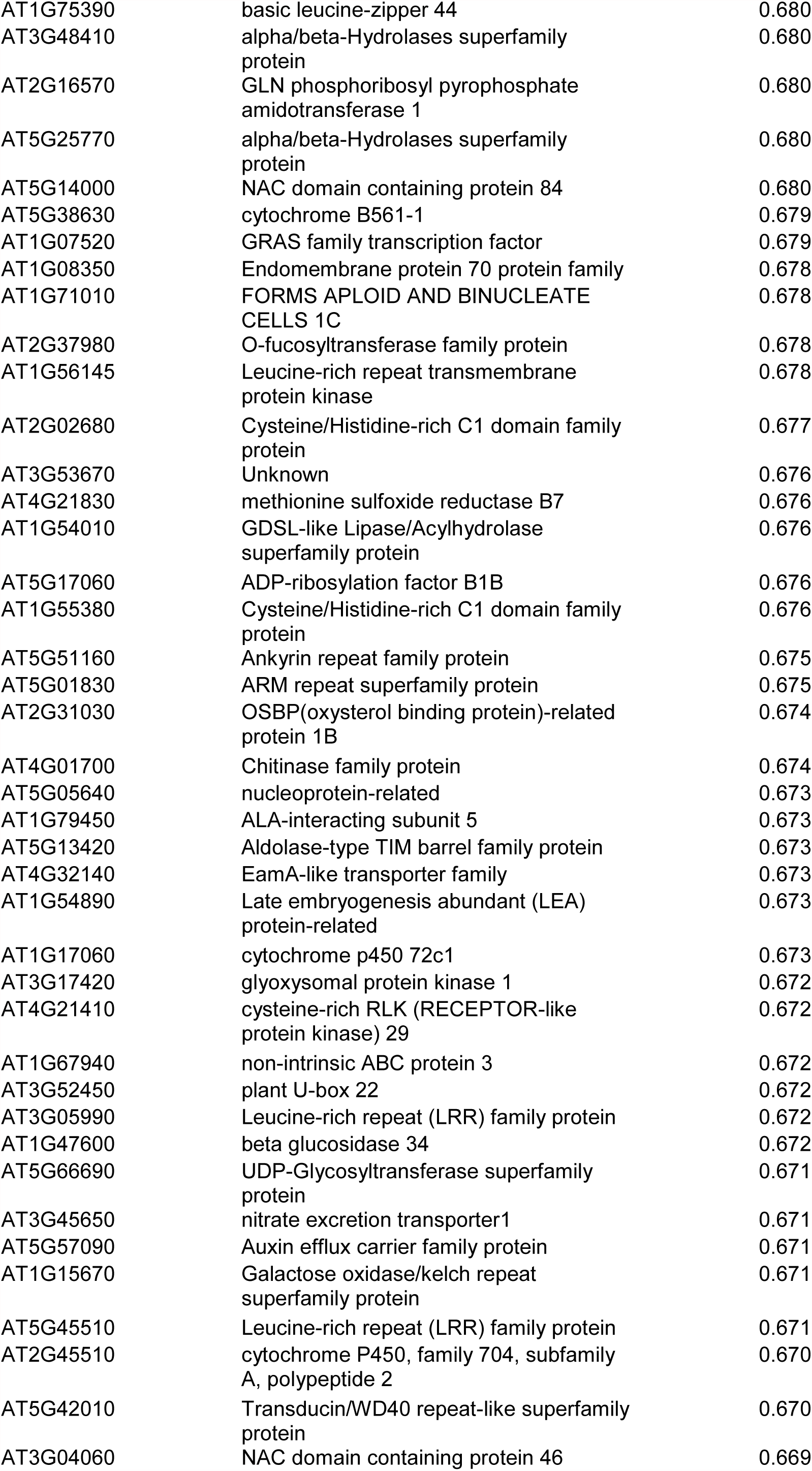
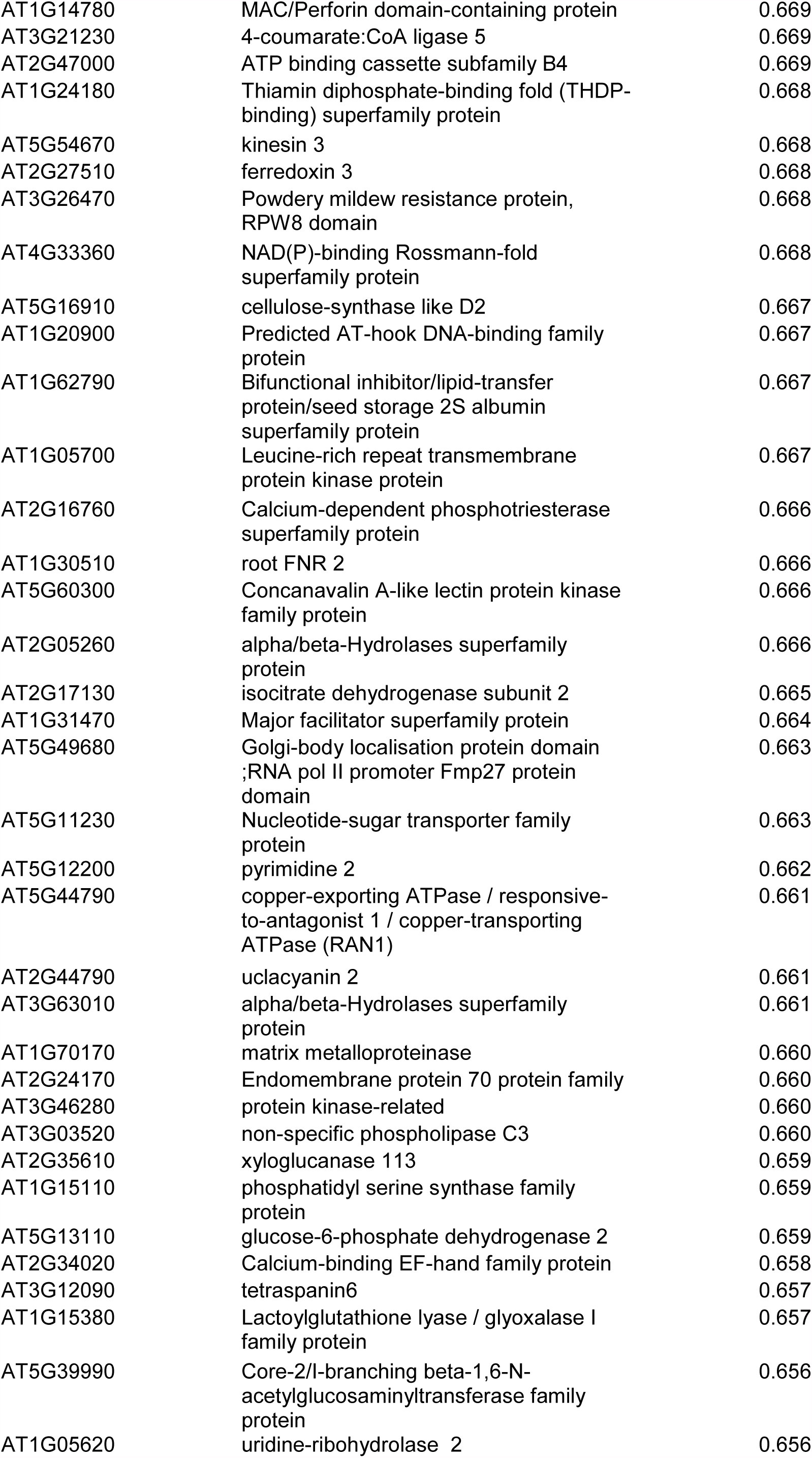
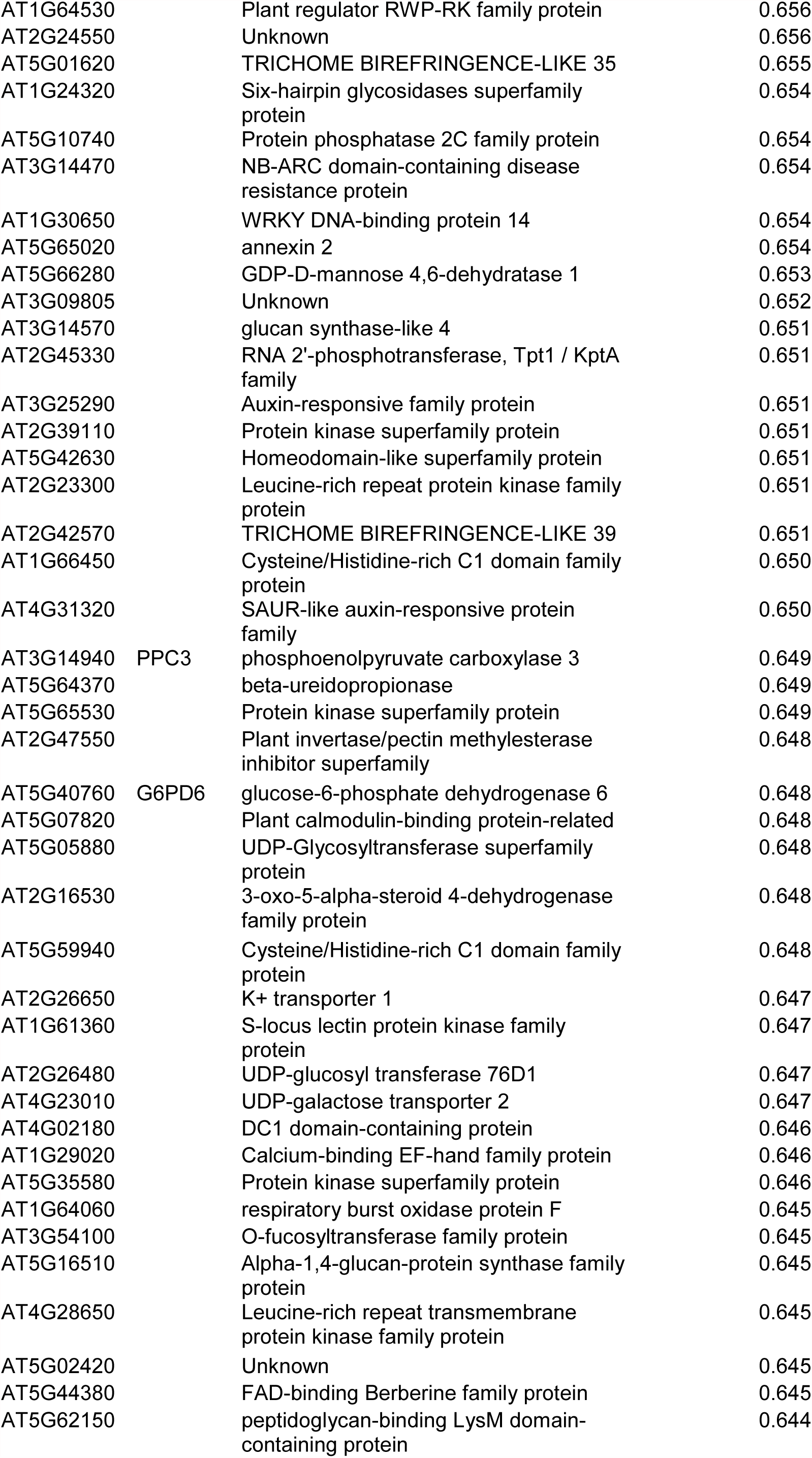
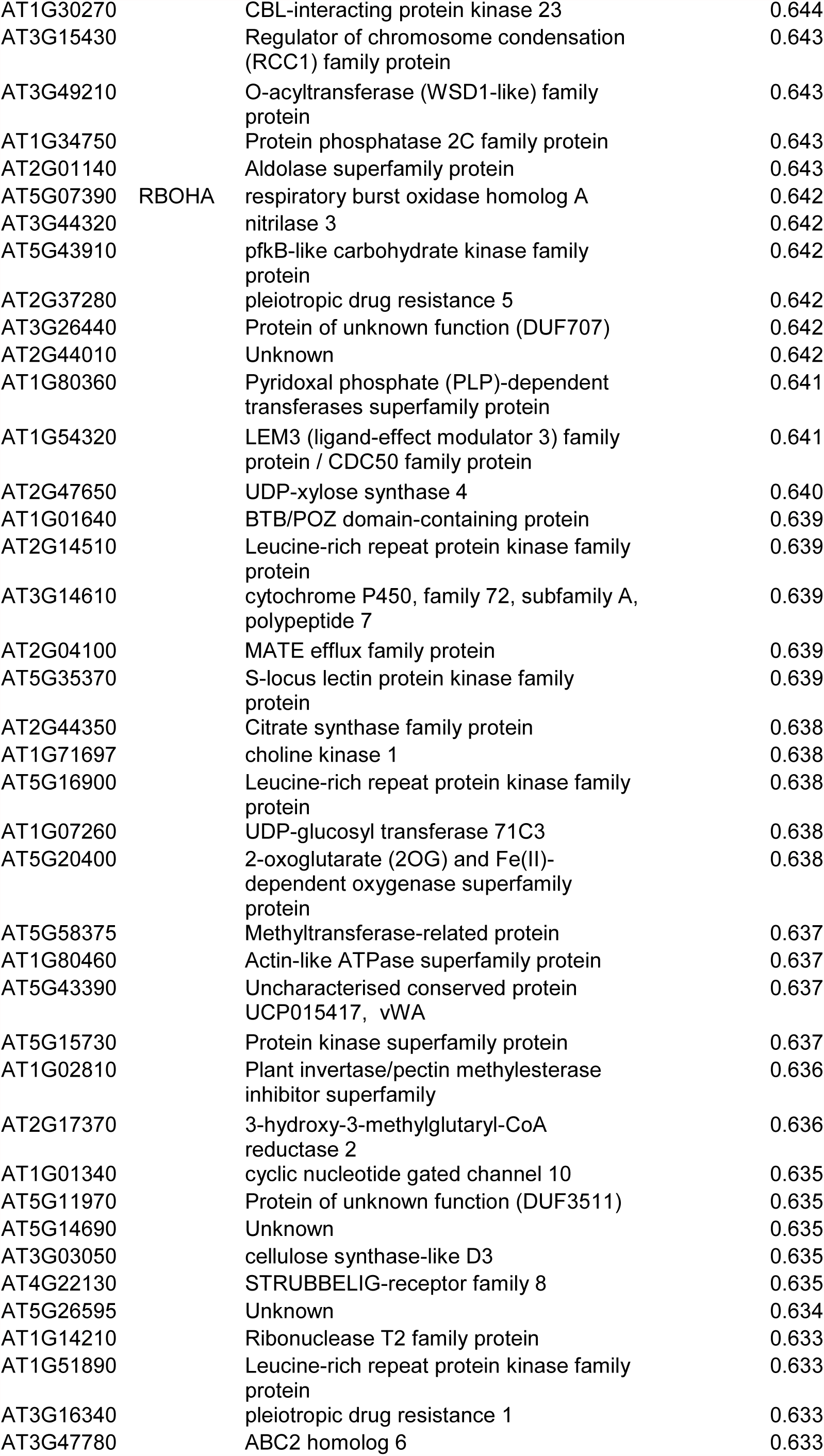
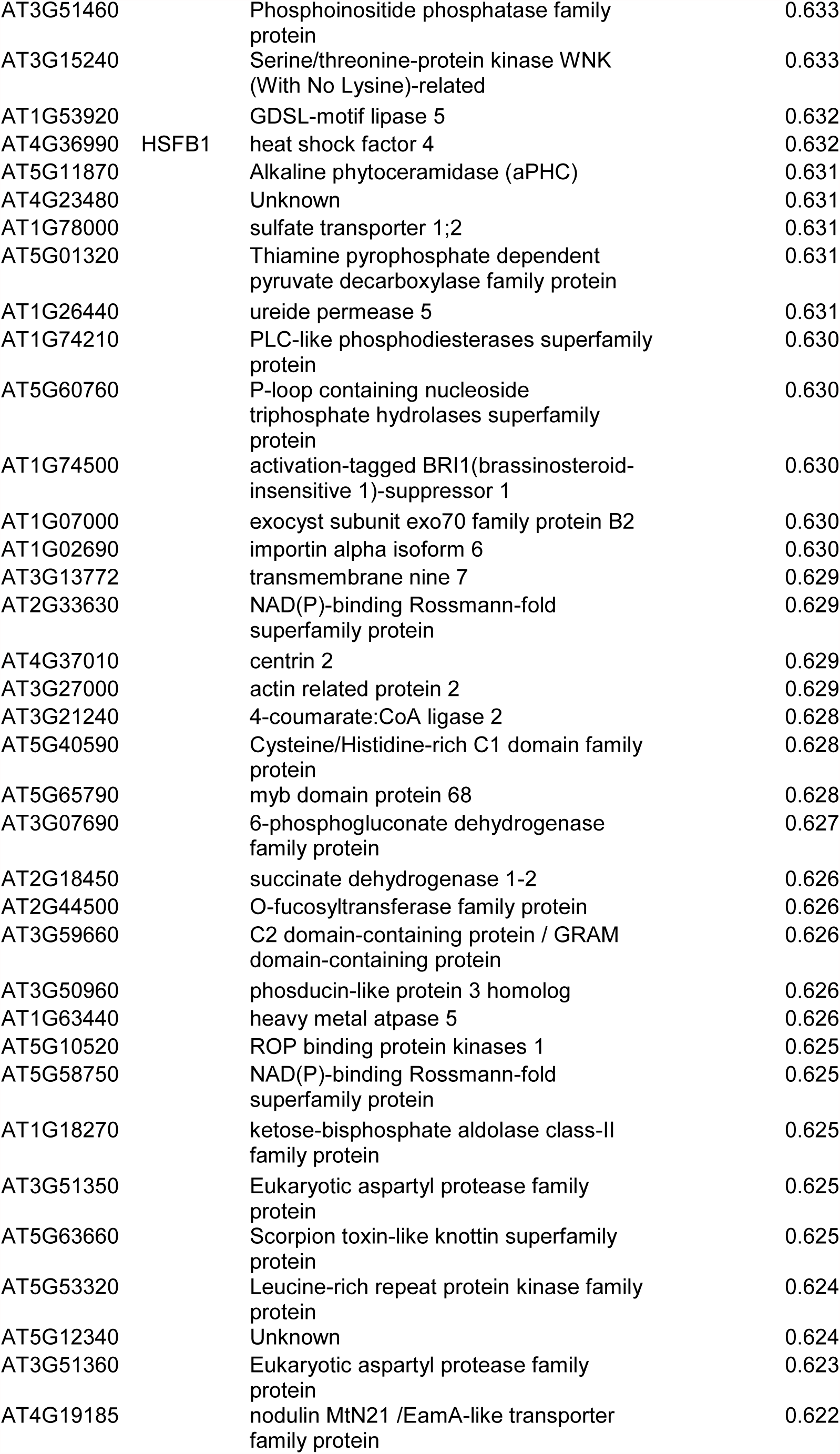
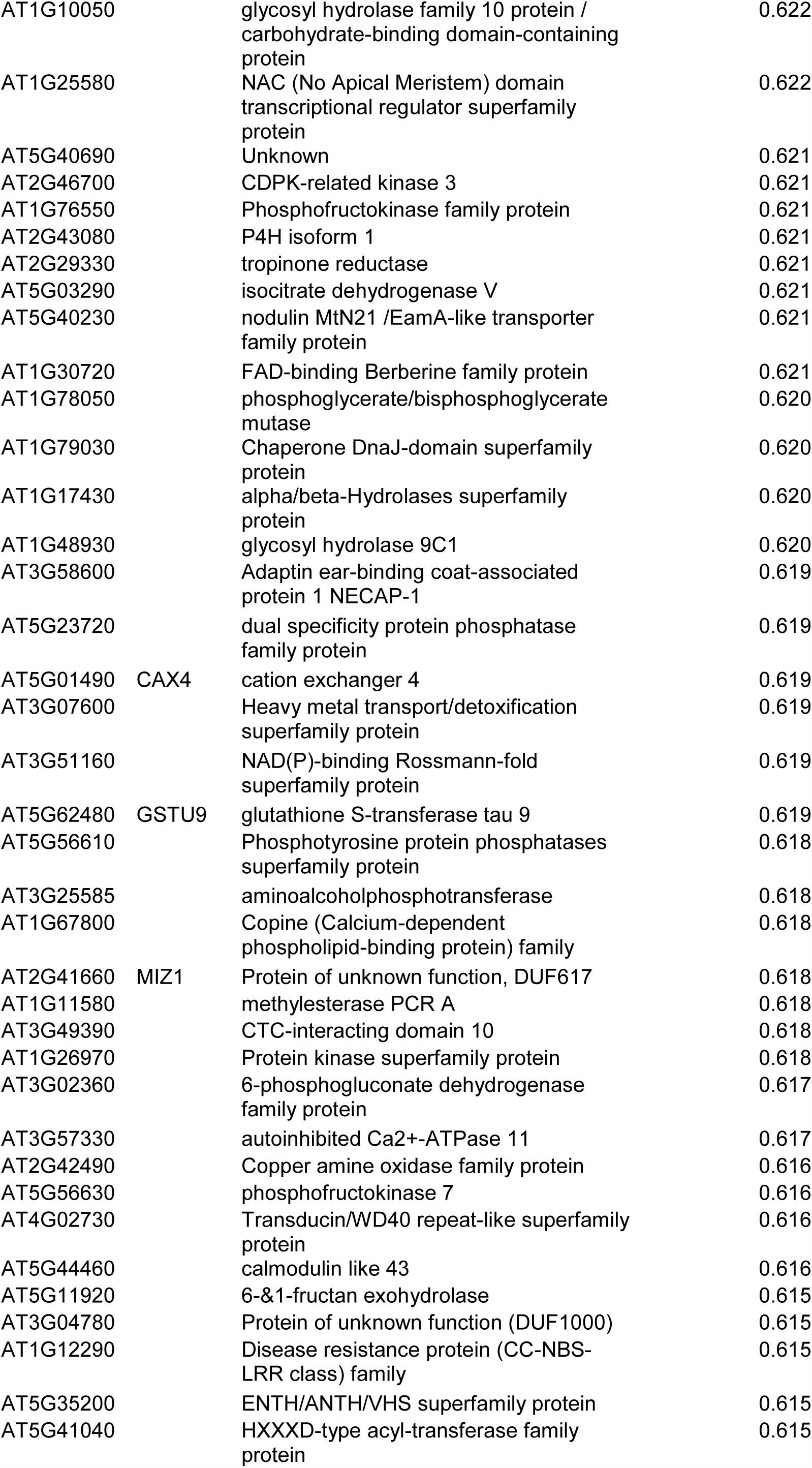
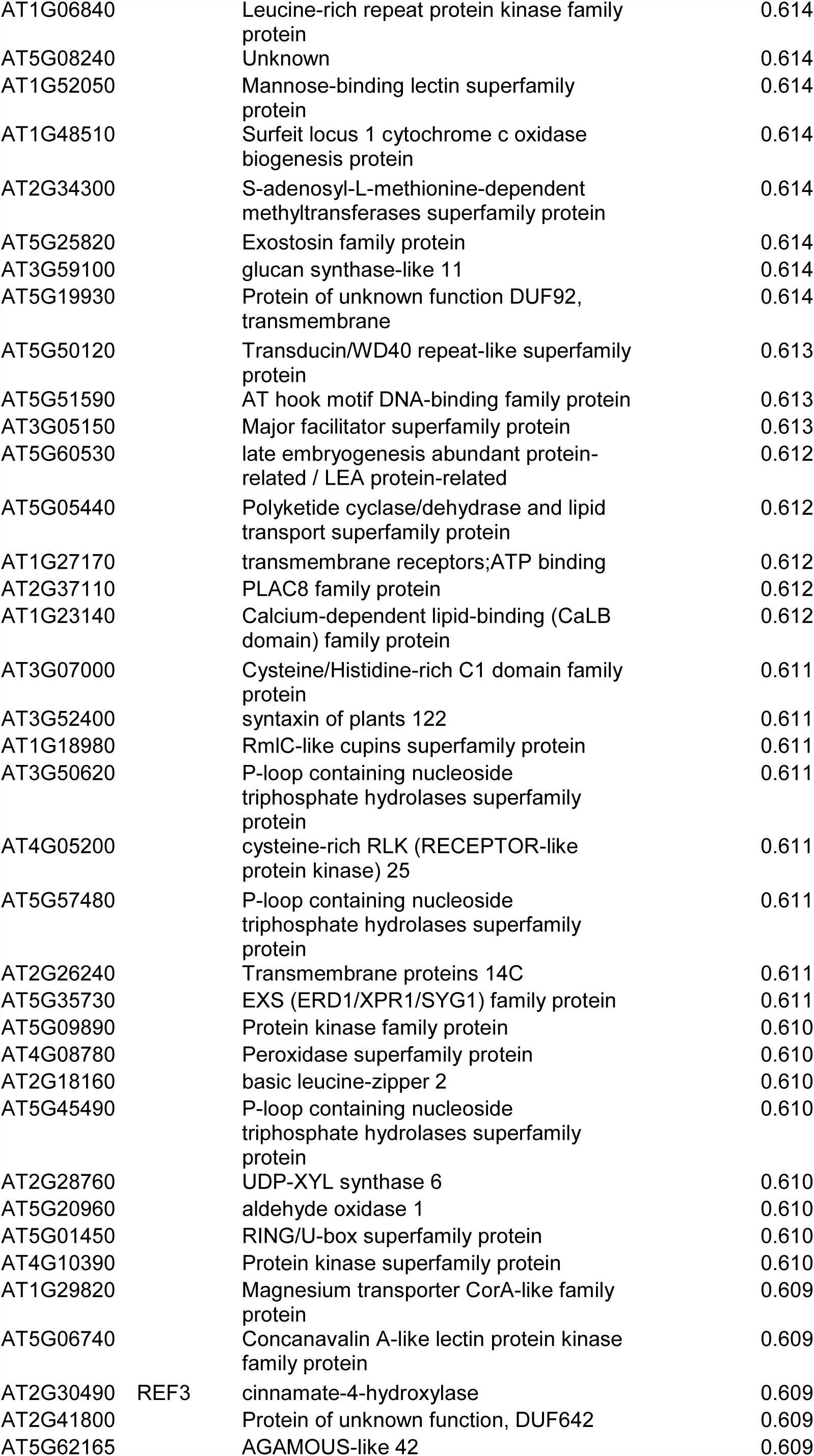
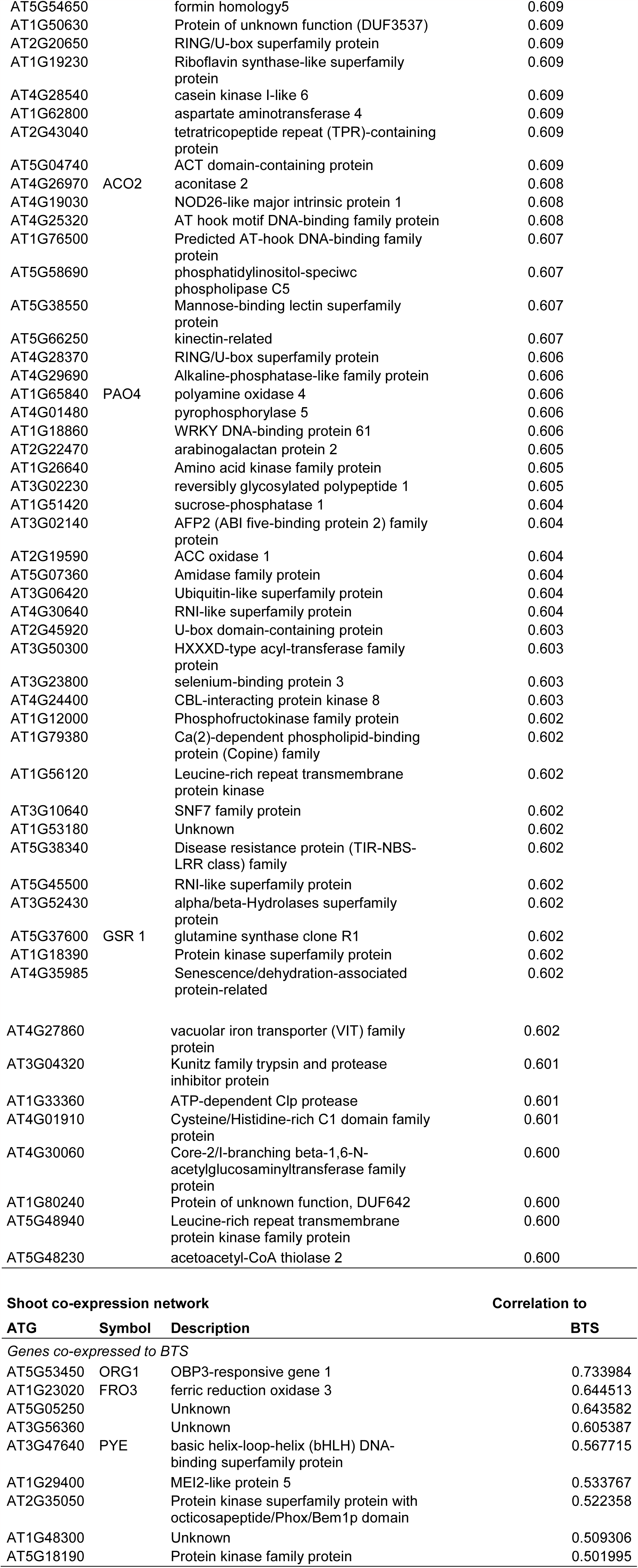
ATG numbers and gene description of genes appearing in the correlation networks shown in Supplemental Figure S1

**Supplementary Table S2.** ATG numbers and gene description of genes appearing in the correlation networks shown in Figure 1.

| ***BTSL1 and BTSL2 root ferrome network*** |  |  | ***Correlation to*** |  |  |
| --- | --- | --- | --- | --- | --- |
| **ATG** | **Symbol** | **Description** | **BTSL1** | **BTSL2** | **BTS** |
| AT1G74770 | BTSL1 | BRUTUS-Like 1, zinc ion binding |  |  |  |
| AT1G18910 | BTSL2 | BRUTUS-Like 2, zinc ion binding |  |  |  |
| AT4G19690 | IRT1 | iron-regulated transporter 1 | 0.687 |  |  |
| AT3G56980 | bHLH39 | basic helix-loop-helix (bHLH) DNA-binding superfamily protein | 0.607 |  |  |
| AT3G07720 | Kelch | Galactose oxidase/kelch repeat superfamily protein | 0.600 |  |  |
| AT3G53480 | PDR9 | pleiotropic drug resistance 9 |  | 0.693 |  |
| AT5G35580 |  | Unknown |  | 0.646 |  |
| AT3G21240 | AT4CL2 | 4-coumarate:CoA ligase 2 |  | 0.628 |  |
| AT5G03570 | IREG2 | iron regulated 2 | 0.765 | 0.711 |  |
| AT1G05700 |  | Leucine-rich repeat transmembrane protein kinase protein | 0.742 | 0.667 |  |
| AT3G13610 | F6H1 | 2-oxoglutarate (2OG) and Fe(II)-dependent oxygenase superfamily protein | 0.702 | 0.793 |  |
| AT3G60330 | HA7 | H(+)-ATPase 7 | 0.675 | 0.719 |  |
| AT3G58810 | MTPA2 | metal tolerance protein A2 | 0.664 | 0.643 |  |
| AT2G29750 |  | Unknown | 0.654 | 0.684 |  |
| AT3G61410 |  | Unknown | 0.649 | 0.632 |  |
| AT2G28160 | FIT | FER-like regulator of iron uptake | 0.649 | 0.748 |  |
| AT2G32270 | ZIP3 | zinc transporter 3 precursor | 0.643 | 0.665 |  |
| AT3G18560 |  | Unknown | 0.626 | 0.659 |  |
| ***BTS shoot ferrome network*** |  |  | ***Correlation to*** |  |  |
| **ATG** | **Symbol** | **Description** | **BTSL1** | **BTSL2** | **BTS** |
| AT3G18290 | BTS | BRUTUS, zinc finger protein-related |  |  |  |
| AT5G53450 | ORG1 | OBP3-responsive gene 1 |  |  | 0.734 |
| AT1G23020 | FRO3 | ferric reduction oxidase 3 |  |  | 0.645 |
| AT5G05250 |  | Unknown |  |  | 0.644 |
| AT3G47640 | PYE | basic helix-loop-helix (bHLH) DNA-binding superfamily protein |  |  | 0.568 |

**Supplementary Table S3.**
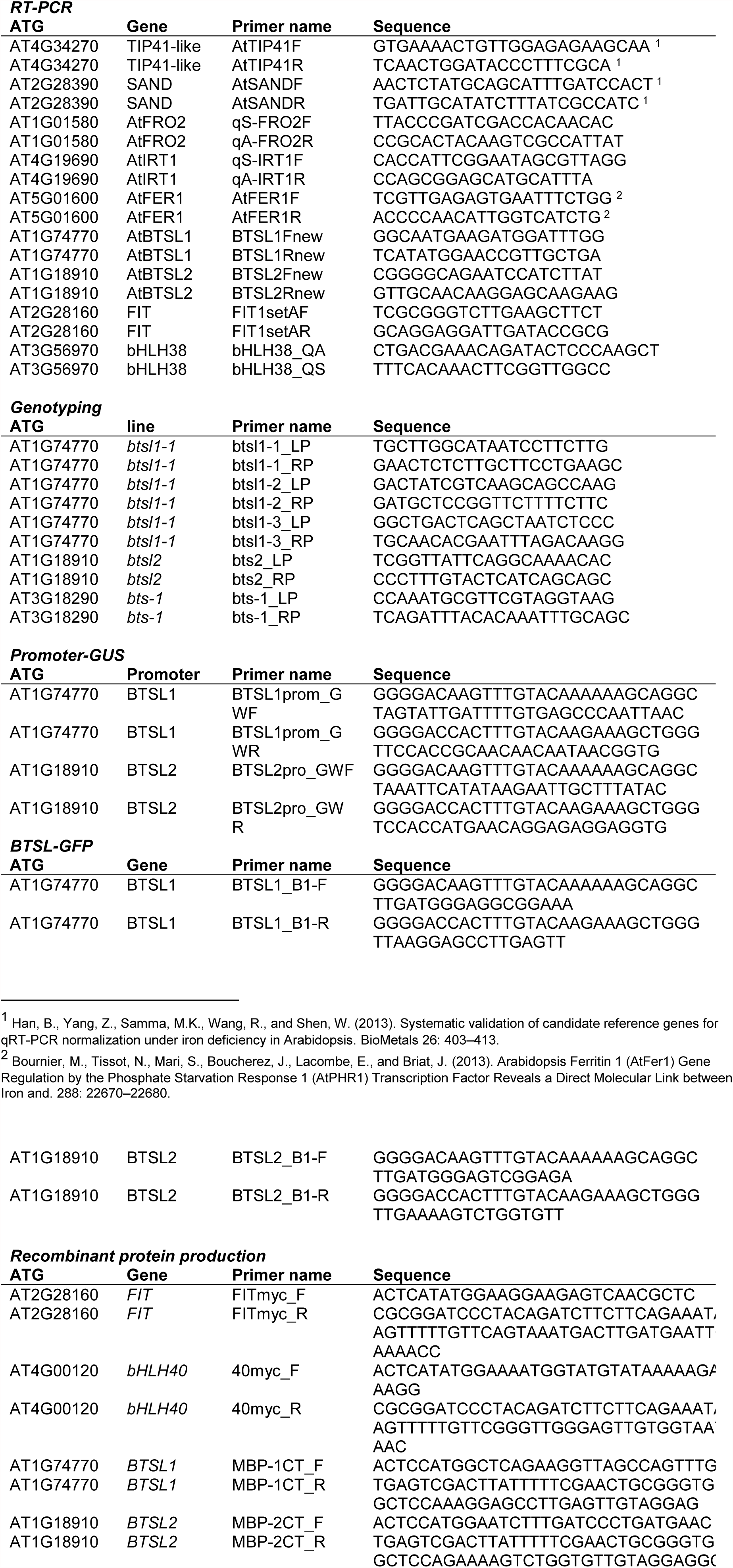
List of primers.

